# Distinct neural progenitor pools in the ventral telencephalon generate diversity in striatal spiny projection neurons

**DOI:** 10.1101/770057

**Authors:** Fran van Heusden, Anežka Macey-Dare, Rohan N. Krajeski, Andrew Sharott, Tommas Jan Ellender

**Affiliations:** Department of Pharmacology, OX1 3QT, Oxford; MRC BNDU, OX1 3TH, Oxford

**Keywords:** basal ganglia, striatum, development, embryonic neural progenitors, spiny projection neurons, apical intermediate progenitors, neural circuits

## Abstract

Heterogeneous populations of neural progenitors in the embryonic lateral ganglionic eminence (LGE) generate all GABAergic spiny projection neurons (SPNs) found in the striatum. Here we investigate how this diversity in neural progenitors relates to diversity of adult striatal neurons and circuits. Using a combination of *in utero* electroporation to fluorescently pulse-label striatal neural progenitors in the LGE, brain slice electrophysiology, electrical and optogenetic circuit mapping and immunohistochemistry, we characterise a population of neural progenitors enriched for apical intermediate progenitors (aIPs) and a distinct population of other progenitors (OPs) and their neural offspring. We find that neural progenitor origin has subtle but significant effects on the properties of striatal SPNs. Although aIP and OP progenitors can both generate D1-expressing direct pathway as well as D2-expressing indirect pathway SPNs found intermingled in the striatum, the aIP derived SPNs are found in more medial aspects of the striatum, exhibit more complex dendritic arbors with higher spine density and differentially sample cortical input. Moreover, optogenetic circuit mapping of the aIP derived neurons show that they further integrate within striatal circuits and innervate both local D1 and D2 SPNs. These results show that it is possible to fluorescently pulse-label distinct neural progenitor pools within the LGE and provide the first evidence that neural progenitor heterogeneity can contribute to the diversity of striatal SPNs.

## Introduction

A fundamental question in neuroscience is how neuronal cell types and neural circuits arise and what critically guides their development. Recent studies in the dorsal telencephalon or pallium have highlighted important and distinct roles for embryonic neural progenitors, at both the level of single neural progenitors (Yu *et al.*, 2009; Yu *et al.*, 2012; Cadwell *et al.*, 2019) and distinct pools of neural progenitors (Tyler *et al.*, 2015; Ellender *et al.*, 2018), in shaping neuronal identity and synaptic connectivity. However, much less is known about how embryonic neural progenitors in the ventral telencephalon or subpallium contribute to the cellular diversity and neural circuitry found within ventral brain structures. Neural progenitors in the ventral portions of the embryonic brain can be found in the ganglionic eminences – transitory structures that generate most interneurons of the brain and the spiny projection neurons (SPNs) of the striatum (Wonders & Anderson, 2006; Wamsley & Fishell, 2017). In this study we focused on the neural progenitors found within the lateral ganglionic eminence (LGE) (Graybiel & Ragsdale, 1978; Olsson *et al.*, 1998; Mason *et al.*, 2005; Pilz *et al.*, 2013; Kelly *et al.*, 2018), which give rise to the striatal SPNs, and include radial glial cells, basal radial glial cells, subapical progenitors, basal progenitors and short neural precursors, amongst others (Olsson *et al.*, 1998; Stenman *et al.*, 2003; Pilz *et al.*, 2013). Many of these are not unique to the LGE and have previously been characterised in detail in the proliferative zones of the cortex (Noctor *et al.*, 2001; Noctor *et al.*, 2004; Gal *et al.*, 2006; Kowalczyk *et al.*, 2009; Stancik *et al.*, 2010; Shitamukai *et al.*, 2011; Wang *et al.*, 2011; Franco & Muller, 2013; Taverna *et al.*, 2014). However, the relative abundance as well as key proliferative behaviours of many of these progenitors differ between cortex and the LGE (Pilz *et al.*, 2013). Furthermore, it is unknown to what extent these different pools of neural progenitors contribute to the diversity of striatal projection neurons.

To investigate the relationship between neural progenitor pool and striatal neuron diversity, we used *in utero* electroporation to fluorescently pulse-label two pools of actively dividing neural progenitors in the LGE distinguished by their differential expression of the tubulin alpha1 (Tα1) promoter (Gal *et al.*, 2006; Stancik *et al.*, 2010). We find that Tα1-expressing neural progenitors exhibit many of the characteristics of the previously described short neural precursors and subapical progenitors found within the LGE (Pilz *et al.*, 2013), including division in the ventricular zone, lack of basal processes during division and relatively fast cell-cycle kinetics. In contrast, the non-Tα1-expressing neural progenitors tend to retain their basal processes during division and exhibit slower cell-cycle kinetics. Overall, our observations suggest that Tα1-expressing neural progenitors consist of intermediate progenitors found in apical regions of the LGE such as short neural precursors and subapical progenitors (Pilz *et al.*, 2013) and we therefore refer to them collectively as apical intermediate progenitors (aIPs) and the non-Tα1-expressing neural progenitors consist of all other neural progenitors that are actively dividing at the same time and we collectively refer to them as other progenitors (OPs). We find that aIP and OP neural progenitors mainly generate striatal neurons, with all the hallmarks of GABAergic spiny projection neurons (SPNs) including expression of neurochemical markers such as CTIP2, but also generate a small number of neurons found within the olfactory bulb. Using the neurochemical marker PPE we find that both aIP and OP neural progenitors generate D1-expressing direct pathway as well as D2-expressing indirect pathway SPNs. Stereological investigation of progenitor derived striatal neurons shows that aIP derived neurons are on average found in more medial aspects of the striatum. Whole-cell patch-clamp recordings of aIP and OP derived neurons in acute striatal slices reveals that their electrical properties are similar and consistent with those of SPNs. Indeed, subsequent anatomical reconstruction of recorded neurons confirms that they exhibit all the hallmarks of SPNs, including radially oriented dendrites with large numbers of spines. Interestingly, we find differences with aIP derived SPNs exhibiting a greater local dendritic complexity as well as a higher density of spines. Lastly, using both electrical and optogenetic circuit mapping we find that both aIP and OP derived SPNs integrate within striatal circuits. Firstly, both receive excitatory glutamatergic input from cortex with aIP derived SPNs exhibiting a significantly prolonged response to cortical activation. Secondly, aIP derived SPNs form local GABAergic inhibitory synaptic connections with neighbouring D1 and D2 SPNs.

In conclusion, this study shows it is possible to label distinct pools of neural progenitors in the LGE and suggests that neural progenitor heterogeneity contributes to striatal diversity in that neural progenitor origin has subtle but significant effects on the spatial distribution of striatal neurons, their morphology as well as their sampling of excitatory inputs.

## Results

### The LGE of the ventral telencephalon contains tubulin αlpha1 (Tα1) expressing neural progenitors

To investigate the relationship between the diverse pools of neural progenitors in the LGE and their contribution to striatal neuronal identity and connectivity we performed *in utero* electroporation (IUE) to fluorescently label actively dividing progenitors in the ventricular zone (Stancik *et al.*, 2010). Two DNA constructs were electroporated into the LGE: a Tα1-cre construct in which cre recombinase is under the control of a portion of the Tα1 promoter (Stancik *et al.*, 2010), and a CβA-FLEx reporter construct that incorporates a flexible excision (FLEx) cassette where cre recombination permanently switches expression from TdTomato fluorescent protein to enhanced green fluorescent protein (GFP) (Franco *et al.*, 2012).

We find that 24 hours after IUE the LGE contains both GFP and TdTomato expressing cells, consisting of Tα1-expressing (Tα1^+^) and non-Tα1-expressing (Tα1^-^) neural progenitors and young migrating neurons (Figure 1A). We find that the time of IUE influenced the relative proportion of GFP and TdTomato expressing cells seen 24 hours later, with IUE at E15.5 resulting in a greater number of Tα1^+^/GFP^+^ cells (E12.5: 4.8 ± 4.1% and E15.5: 34.5 ± 3.0%, Mann-Whitney test, p=0.002, n = 7 and 17 embryonic brains, Figure 1B), suggesting that, similar to their cortical counterparts (Stancik *et al.*, 2010; Ellender *et al.*, 2018) Tα1-expressing progenitors form a considerable population of actively proliferating cells during later periods of neurogenesis.

**Figure 1:**
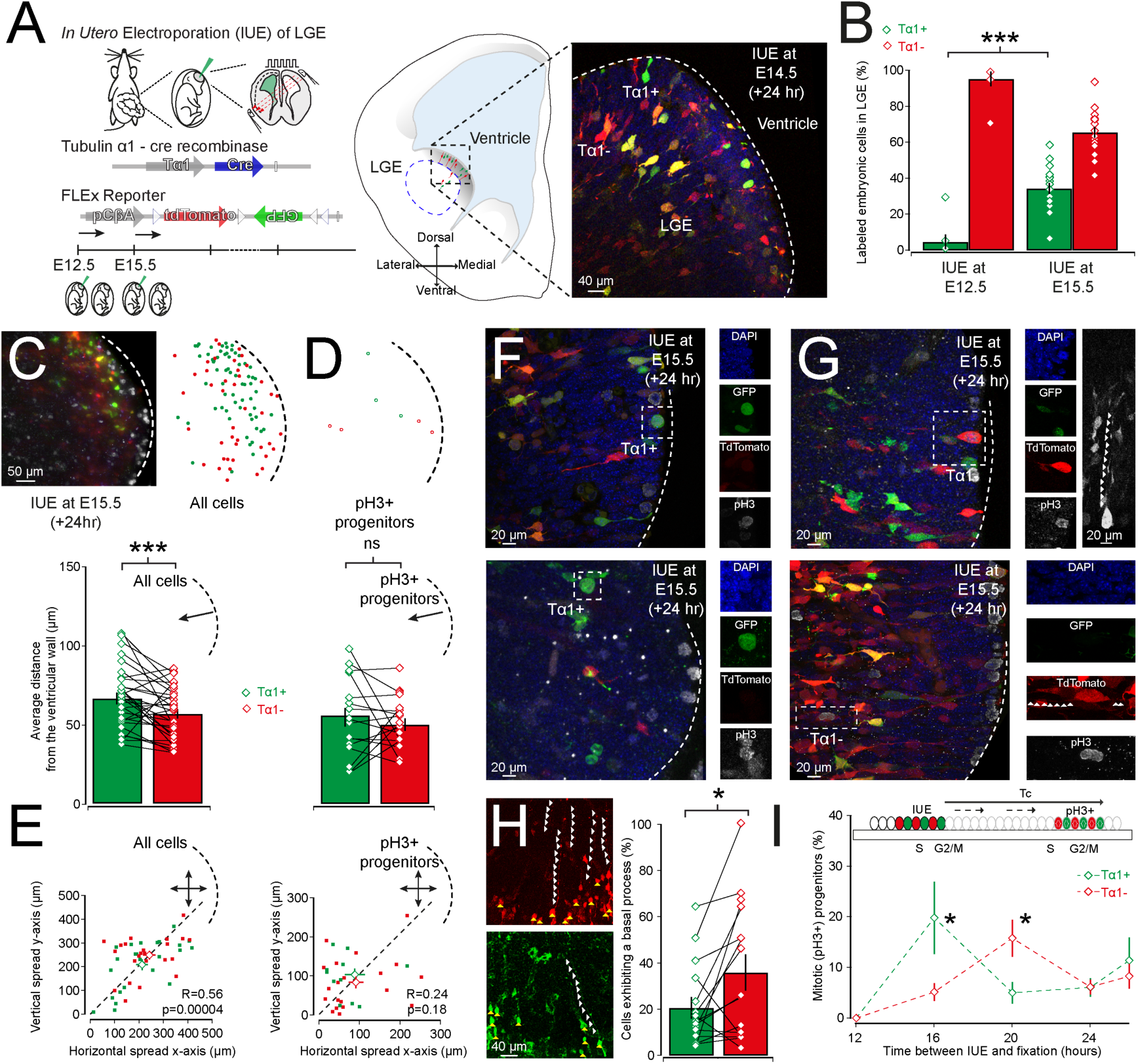
The LGE contains both Tα1-expressing and non-Tα1-expressing neural progenitors. (**A**) *In utero* electroporation (IUE) of Tα1-cre and FLEx reporter plasmids was performed at embryonic day (E)12.5 and E15.5 to label Tα1-positive (Tα1^+^) and Tα1-negative (Tα1^-^) neural progenitors lining the ventricular wall of the LGE with respectively the fluorescent markers GFP or TdTomato (left). 24 hours after IUE fluorescently labeled Tα1^+^/GFP^+^ and Tα1^-^/TdTomato^+^ neural progenitors and young neurons can be found in the LGE (right). Yellow cells were were assumed to have undergone recombination relatively recently and counted as GFP^+^. (**B**) Quantification of Tα1^+^/GFP^+^ and Tα1^-^/TdTomato^+^ cells reveals that the Tα1^+^/GFP^+^ cells form a substantial population during late stages of embryogenesis. (**C**) The distance from the ventricle of Tα1^+^/GFP^+^ and Tα1^-^/TdTomato^+^ cells 24 hours after IUE differs in that Tα1^+^/GFP^+^ cells are located significantly further away from the ventricular wall. (**D**) A difference in distance from the ventricle is not seen for actively dividing Tα1^+^/GFP^+^ and Tα1^-^/TdTomato^+^ cells. (**E**) Horizontal and vertical migration of all labeled cells exhibits a linear correlation (left) whereas this is not the case for actively dividing progenitors (right). (**F**) Actively dividing Tα1^+^/GFP^+^ progenitors exhibiting a short morphology and lacking a basal endfoot can be found dividing at the ventricular wall (top) and at a small distance from ventricle (bottom), corresponding to respectively a short neural precursor and subapical progenitor. (**G**) In contrast, dividing Tα1^-^/TdTomato^+^ cells predominantly retain a basal process during division and can be found dividing at both the ventricular wall (top) and at a distance from the ventricle (bottom) corresponding to respectively a radial glial cell and bipolar radial glial cell. (**H**) Quantification of cells in the proliferative zones of the LGE containing a basal process. Note that significantly more Tα1^-^/TdTomato^+^ cells retain a basal process. (**I**) Estimation of cell cycle duration using embryonic tissue fixed at variable delays after IUE and labeling for the mitotic marker phospho-histone H3 (pH3). A rapid increase in the number of neural progenitors that were positive of the marker pH3 was taken as a return to the mitotic (‘M’) phase for that population of neural progenitors and a proxy for the cell cycle duration (‘Tc’).

Using IUE at E15.5 we next investigated the spatial distribution of Tα1^+^/GFP^+^ and Tα1^-^/TdTomato^+^ cells within the LGE to further characterize these cells and gain insight into their proliferative behaviour. Firstly, we assessed the location of all Tα1^+^/GFP^+^ and Tα1^-^/TdTomato^+^ cells and dividing progenitors relative to the ventricular wall 24 hours after IUE (Figure 1C and D). Tα1^+^/GFP^+^ and Tα1^-^ /TdTomato^+^ cells can be found interspersed in the proliferative zone of the LGE with the Tα1^+^/GFP^+^ labeled cells on average significantly further away from the ventricular wall than the Tα1^-^/TdTomato^+^ labeled cells (Tα1^+^/GFP^+^: 66.24 ± 3.57 µm and Tα1^-^/TdTomato^+^: 56.45 ± 2.90 µm, paired t-test, p=0.0055, n = 30, Figure 1C). However, when we limited analysis to only actively dividing progenitor cells, labeled with the mitotic marker phospho-histone H3 (pH3), no difference was seen in their distance from the ventricle (Tα1^+^/GFP^+^: 52.28 ± 6.21 µm and Tα1^-^/TdTomato^+^: 48.78 ± 3.99 µm, p=0.586, paired t-test, n = 16 brains, Figure 1D). These results are consistent with both Tα1^+^/GFP^+^ and Tα1^-^/TdTomato^+^ progenitors dividing mainly within the ventricular zone of the LGE (Pilz *et al.*, 2013). Furthermore, our results suggest that 24 hours after IUE Tα1^+^/GFP^+^ cells, likely consisting of young migrating neurons and non-dividing neural progenitors, inhabit increasingly subapical positions within the LGE. Whereas in the cortex young migrating neurons mostly exhibit radial migration (Noctor *et al.*, 2001) newly born neurons in the striatum undergo tangential, radial and other forms of migration (Halliday & Cepko, 1992; Tan & Breen, 1993; Reid & Walsh, 2002; Tinterri *et al.*, 2018). Indeed, when we look at the average dispersion of all labeled cells in a subset of brain slices along two axis perpendicular to the ventricular wall we find a positive linear correlation between the amount of horizontal and vertical spread of labeled cells (R=0.56, p=0.00004, n=26 slices) which is similar for both Tα1^+^/GFP^+^ and Tα1^-^/TdTomato^+^ cells (p>0.05) and is not seen when we limit the analysis to only actively dividing neural progenitors (R=0.24, p=0.18, n=20 slices, Figure 1E). Lastly, the LGE has also been shown to consist of several domains depending on differential expression of certain transcription factors (Stenman *et al.*, 2003; Flames *et al.*, 2007; Xu *et al.*, 2018). When we split the LGE in 4 different domains we find that both Tα1^+^/GFP^+^ and Tα1^-^/TdTomato^+^ cells can be found in all progenitor domains of the LGE (Supplemental Figure 1).

We next investigated the morphology of actively dividing neural progenitors in the LGE. The distinguishing features of short neural precursors in the dorsal and ventral telencephalon are their division at the ventricular wall and their short morphology during division lacking a prominent basal process contacting pial surfaces (Gal *et al.*, 2006; Stancik *et al.*, 2010; Pilz *et al.*, 2013; Ellender *et al.*, 2018). Indeed, when we look at the morphology of labeled neural progenitors we find that many actively dividing Tα1^+^/GFP^+^ progenitors exhibit a short rounded morphology during division and could be found at the ventricular wall (Figure 1F). However, some Tα1^+^/GFP^+^ progenitors divide slightly away from the ventricular wall resembling subapical progenitors (Pilz *et al.*, 2013) (Figure 1F). In contrast, many actively dividing Tα1^-^/TdTomato^+^ progenitors retained their basal process during division and could be found either at or away from the ventricular wall (Figure 1G). This was confirmed quantitatively by assessment of the number of cells in the VZ that retained a basal process. Overall, we find that Tα1^+^/GFP^+^ cells often lacked a basal process in contrast to Tα1^-^/TdTomato^+^ cells that retained theirs (Tα1^+^/GFP^+^: 20.6 ± 4.7% and Tα1^-^/TdTomato^+^: 35.9 ± 7.8%, p=0.03, Wilcoxon signed rank test, n = 15 brains, Figure 1H).

A further distinguishing characteristic of both short neural precursors and subapical progenitors found in the LGE is their relatively fast cell cycle kinetics (Pilz *et al.*, 2013). To investigate whether the Tα1^+^/GFP^+^ progenitor population exhibits fast cell cycle durations we performed IUE at E15.5 and subsequently fixed tissue at varying delays ranging from 12 to 26 hours and labeled dividing neural progenitors with the mitotic marker pH3 (Stancik *et al.*, 2010). Consistent with the fast cell cycle kinetics described for the short neural precursors and subapical progenitors in the LGE we find that the Tα1^+^/GFP^+^ progenitors return to G1/S phase faster that the Tα1^-^ /TdTomato^+^ progenitors. Indeed, after a 16 hour delay Tα1^+^/GFP^+^ progenitors are the dominant population of pH3 expressing cells (Tα1^+^/GFP^+^: 19.85 ± 7.15% and Tα1^-^ /TdTomato^+^: 5.18 ± 1.72%, t-test, p = 0.025, n = 6 brains) whereas after a 20 hour delay it is the Tα1^-^/TdTomato^+^ progenitors which form the dominant population of pH3 expressing cells (Tα1^+^/GFP^+^: 4.98 ± 2.11% and Tα1^-^/TdTomato^+^: 15.81 ± 3.63%, t-test, p = 0.036, n = 8 brains, Figure 1I).

In conclusion, our results suggest that the LGE in the ventral telencephalon contains a population of Tα1-expressing neural progenitors that share many characteristics with short neural precursors and subapical progenitors previously described (Pilz *et al.*, 2013), including a ventricular location of division, a short round morphology during division and fast cell-cycle kinetics. In contrast we are also able to label a population of neural progenitor that does not express Tα1 that share many characteristics with radial glial cells (Pilz *et al.*, 2013) and which tend to retain its basal process and exhibits slower cell cycle kinetics.

### Both progenitor pools can generate striatal D1 and D2 spiny projection neurons

Our *in utero* electroporation labeling experiments suggest that Tα1 expression can distinguish between two pools of neural progenitor in the LGE. The Tα1-expressing neural progenitors exhibit many of the characteristics of described short neural precursors and subapical progenitors whereas, in contrast, the non-Tα1-expressing neural progenitors tend to share many characteristics with the described radial glial cells within the LGE (Pilz *et al.*, 2013). Overall, our observations would suggest the pool of Tα1-expressing neural progenitors is enriched for apically dividing intermediate progenitors and therefore we will refer to them collectively as apical intermediate progenitors (aIPs) and the neurons they generate as aIP derived and the non-Tα1-expressing neural progenitors consist of all other neural progenitors that are actively dividing at the same time and we will refer to them collectively as other progenitors (OPs) and the neurons they generate as OP derived.

When we let the electroporated embryos develop postnatally we observe many GFP and TdTomato expressing neurons spread throughout the striatum (Figure 2B). The GFP expressing neurons are generated from the population of aIP neural progenitors and are referred to as aIP derived and the TdTomato expressing neurons from the population of OP neural progenitors and are referred to as OP derived (Figure 2B). At higher magnifications both pools of labeled neurons exhibit the morphology of striatal spiny projection neurons (SPNs) including dendrites densely covered with spines (Figure 2B). The relative proportion of aIP and OP derived neurons seen postnatally is very similar to the ratio of labeled embryonic neural progenitors found 24 hours after IUE at E15.5, with a significantly larger proportion of OP derived neurons (aIP derived: 36.7 ± 5.5% and OP derived: 63.3 ± 5.5%, p=0.0017, paired t-test, n = 16 brains, Figure 2C). We next investigated the spatial distribution of aIP and OP derived neurons in rostral, central and caudal sections of the striatum in a total of 15 electroporated brains (Supplemental Figure 2). In general, across all sections we find that the normalised position (see **Methods**) of all labeled SPNs was biased to the medial aspects (Wilcoxon Sign Rank, p = 0.000005, n = 15 mice / 23 sections, Figure 2D), but not to either dorsal or ventral aspects of the striatum (Wilcoxon Sign Rank, p > 0.05). We find that within individual sections the average location of aIP derived and OP derived neurons differ suggesting that they might exhibit different migration patterns (Halliday & Cepko, 1992; Tan & Breen, 1993; Reid & Walsh, 2002; Tinterri *et al.*, 2018). Although this likely is not captured fully in coronal slices, we nonetheless find that aIP derived neurons were approximately twice as likely to be statistically biased (Wilcoxon Sign rank, p < 0.016) towards the medial aspect of striatum (73% of sections) than OP derived neurons (37% of sections) with the strongest medial bias seen in the central and caudal planes (Supplemental Figure 2).

**Figure 2:**
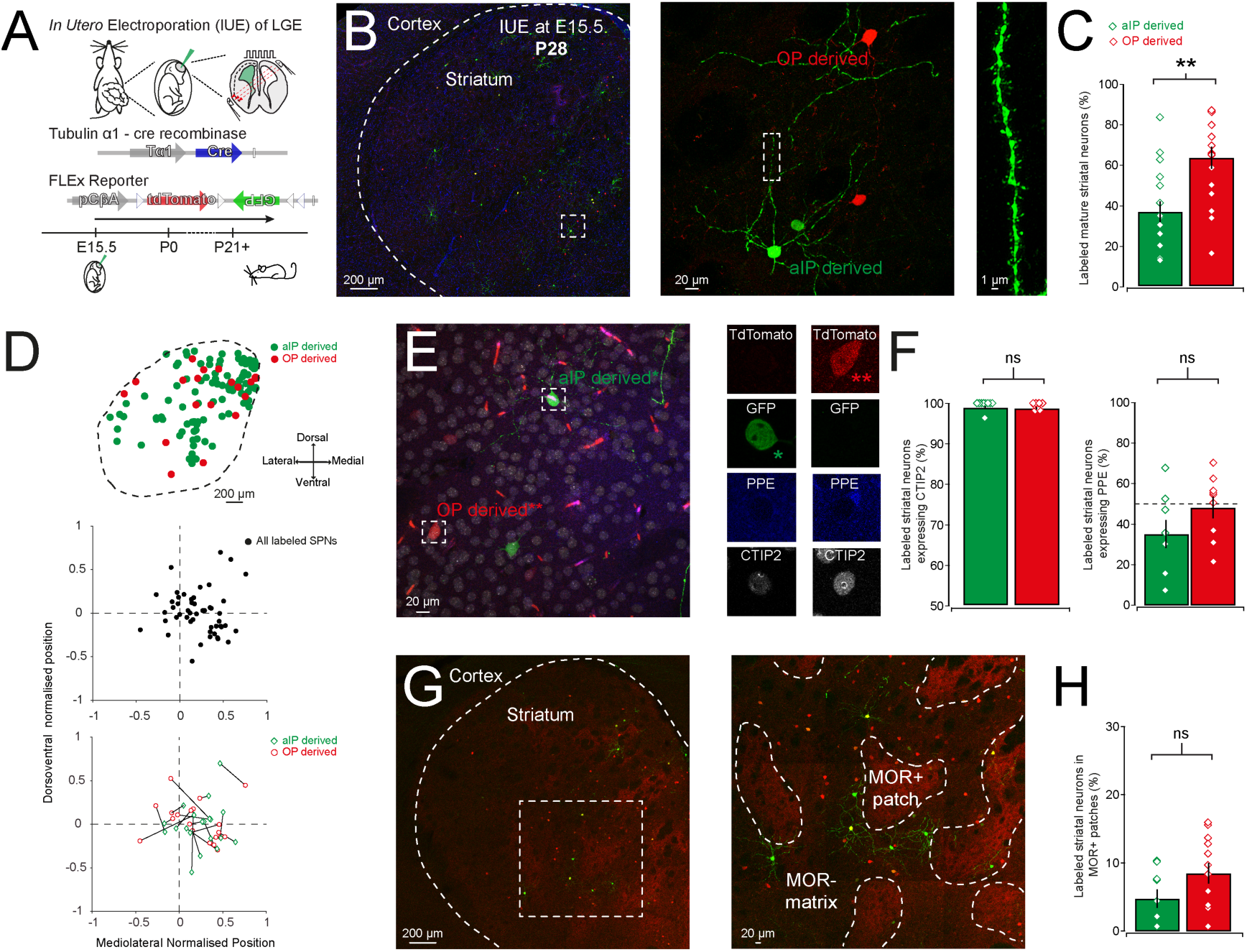
Both aIP and OP neural progenitors in the LGE can generate D1 and D2 striatal spiny projection neurons. (**A**) Mice that underwent IUE at E15.5 with Tα1-cre and FLEx reporter plasmids were left to grow up untill young adulthood (postnatal day 21 and over). (**B**) The striatum of these mice contain both GFP^+^ and TdTomato^+^ neurons derived from respectively a pool of apical intermediate progenitors (aIPs) or other progenitors (OPs). The adult neurons derived from both progenitor pools display the radial morphology and spine studded dendrites (right) characteristic of striatal spiny projection neurons (SPNs). (**C**) Quantification of the relative proportion of aIP and OP derived striatal neurons reveals a similar distribution as the respective embryonic neural progenitors at E15.5. Note the significantly larger proportion of OP derived neurons. (**D**) Example diagram of location of aIP and OP derived neurons in a central section of the striatum (top). Normalised mediolateral and dorsoventral positions of all labeled neurons across all counted sections reveals that labeled neurons inhabit more medial aspects of the striatum (middle). Normalised mediolateral and dorsoventral positions of aIP and OP derived neurons within individual sections, linked with lines, shows that in many sections they reside in different striatal regions (bottom). (**E**) An example image of striatum containing aIP and OP derived neurons and labeled for the SPN marker CTIP2 (white) and the D2-SPN marker PPE (blue). The red streaks result from auto-fluorescence of blood vessels. (**F**) Virtually all labeled neurons are positive for the SPN marker CTIP2 and both aIP and OP derived neurons consist of PPE positive (D2) and PPE negative (putative D1) SPNs. (**G**) Labeling for the µ-opioid receptor (MOR) reveals that aIP and OP derived neurons can be found in both MOR^+^ patch and MOR^-^ matrix compartments. (**H**) Quantification of aIP and OP derived neurons labeled for MOR reveals both can be found within MOR^+^ patches in equal numbers.

We next investigated which types of striatal SPN were labeled. Combining immunocytochemistry for the marker for spiny projection neurons CTIP2 (Arlotta *et al.*, 2008) and the marker for D2 SPNs PPE (Gerfen *et al.*, 1990) we investigated the degree of co-localization of these markers with our aIP and OP derived striatal neurons (Figure 2E). We find that nearly all labeled neurons co-localize with CTIP2 (aIP derived/CTIP2^+^: 99.6 ± 0.4% and OP derived/CTIP2^+^: 99.4 ± 0.3%, both n = 9, Figure 2F) suggesting the labeled neurons are SPNs. Secondly, labeling for the D2 SPN marker PPE reveals that both progenitor pools can generate both PPE^-^ D1-expressing direct pathway SPNs and PPE^+^ D2-expressing indirect pathway SPNs (aIP derived/PPE^+^: 35.2 ± 7.0% and OP derived/PPE^+^: 48.2 ± 5.3%, n = 9) with a trend for the aIP progenitor pool to generate more PPE^-^ D1 SPNs (vs. 50%, p<0.071, one-sample t-test, n = 8). Lastly, we asked whether SPNs derived from these distinct progenitor pools end up in different compartments of the striatum. We used antibody staining against the µ-opioid receptor (MOR) to delineate the MOR-rich patches and the MOR-poor matrix compartments of the striatum (Figure 2G). We find that on average only 6% of labeled neurons can be found in the striatal patches with the remainder found in the matrix with no difference between aIP and OP derived neurons (aIP/MOR^+^: 4.8 ± 1.3% and OP/MOR^+^: 7.8 ± 1.6 %, n = 12, Figure 2H). As different embryonic stages have been shown to differently contribute to the formation of the striatal neurons and compartments (Brand & Rakic, 1979; Graybiel & Hickey, 1982; Marchand & Lajoie, 1986; van der Kooy & Fishell, 1987; Fishell *et al.*, 1990; Newman *et al.*, 2015; Kelly *et al.*, 2018; Tinterri *et al.*, 2018) we repeated these experiment using IUE at E12.5 (Supplemental Figure 3). We find broadly similar results in that the proportion of labeled adult neurons reflect the ratio of embryonic progenitors with less aIP derived neurons labeled during this earlier stage of neurogenesis (aIP derived: 6.6 ± 4.0% and OP derived: 93.4 ± 4.0%, n = 10, Supplemental Figure 3B), virtually all labeled neurons express the SPN marker CTIP2 (aIP derived/CTIP2^+^: 100.0 ± 0.0% and OP derived/CTIP2^+^: 99.2 ± 0.5%, n = 6, Supplemental Figure 3C) and both progenitor populations can generate putative D1-expressing direct pathway SPNs and D2-expressing indirect pathway SPNs (aIP derived/PPE^+^: 19.2 ± 3.6 % and OP-derived/PPE^+^: 34.0 ± 6.0% n = 8, Supplemental Figure 3D). However, labeling at this younger embryonic age seems to generate fewer indirect pathway D2 SPNs (E12.5 vs. E15.5, Mann-Whitney test, p = 0.037; n = 8 and 12) and slightly more labeled neurons in MOR^+^ patches (E12.5: 12.4 ± 2.4% and E15.5: 7.0 ± 1.3% in MOR^+^ patches; Mann-Whitney test, p = 0.170, n = 4 and 12, Supplemental Figure 3E). Lastly, the LGE has been shown to not only generate striatal SPNs, but has also been shown to generate interneurons found in the olfactory bulb (Stenman *et al.*, 2003). Therefore, we investigated whether we could find labeled neurons also in the olfactory bulb in a small subset of mice that had undergone IUE at E15.5. We did find small numbers of labeled neurons in sections of the olfactory bulb (∼40 neurons per section) with the majority being OP derived (aIP derived olfactory bulb interneurons: 3.9 ± 0.7% and OP derived olfactory bulb interneurons: 96.1 ± 0.7, n = 5 brains, Supplemental Figure 4).

Overall, these results suggest that aIP and OP neural progenitors in the LGE mainly generate D1 and D2 spiny projection neurons, which can be found intermingled in the striatum as well as in patch and matrix compartments. Furthermore, a small number of olfactory bulb neurons are generated mainly from the population of OP neural progenitors.

### Both aIP and OP derived neurons have electrical and morphological properties consistent with SPNs

So far we have shown that we can pulse-label two different pools of neural progenitors in the LGE and that both pools can generate D1 and D2 striatal SPNs based on their neurochemical profile. We next investigated in more detail whether the labeled neurons exhibit the electrophysiological and morphological properties of SPNs. To do this we performed whole-cell patch-clamp recordings of aIP and OP derived neurons as well as unlabeled neurons in dorsal striatum (Figure 3A). The inclusion of biocytin in the intracellular solution allowed for post-hoc labeling and reconstruction of recorded cells (Figure 3B and C). We investigated a variety of electrophysiological properties, including membrane potential (aIP derived: −80.3 ± 1.2 mV, OP derived: −78.9 ± 0.7 mV and unlabeled: −78.3 ± 1.2 mV; p>0.05, t-test, n = 15, 34 and 18 neurons), input resistance (aIP derived: 88.1 ± 8.4 MΩ, OP derived: 81.0 ± 4.7 MΩ and unlabeled: 80.3 ± 7.5 MΩ; p>0.05, t-test), action potential threshold (aIP derived: −41.6 ± 1.8 mV, OP derived: −39.8 ± 0.9 mV and unlabeled: - 39.5 ± 1.2 mV; p>0.05, t-test) as well as action potential frequency, which all suggested that labeled neurons exhibited electrical properties consistent with those of SPNs. However, we did not observe significant differences in these or other electrophysiological properties between aIP and OP derived neurons (Figure 3D and Table 1).

**Figure 3:**
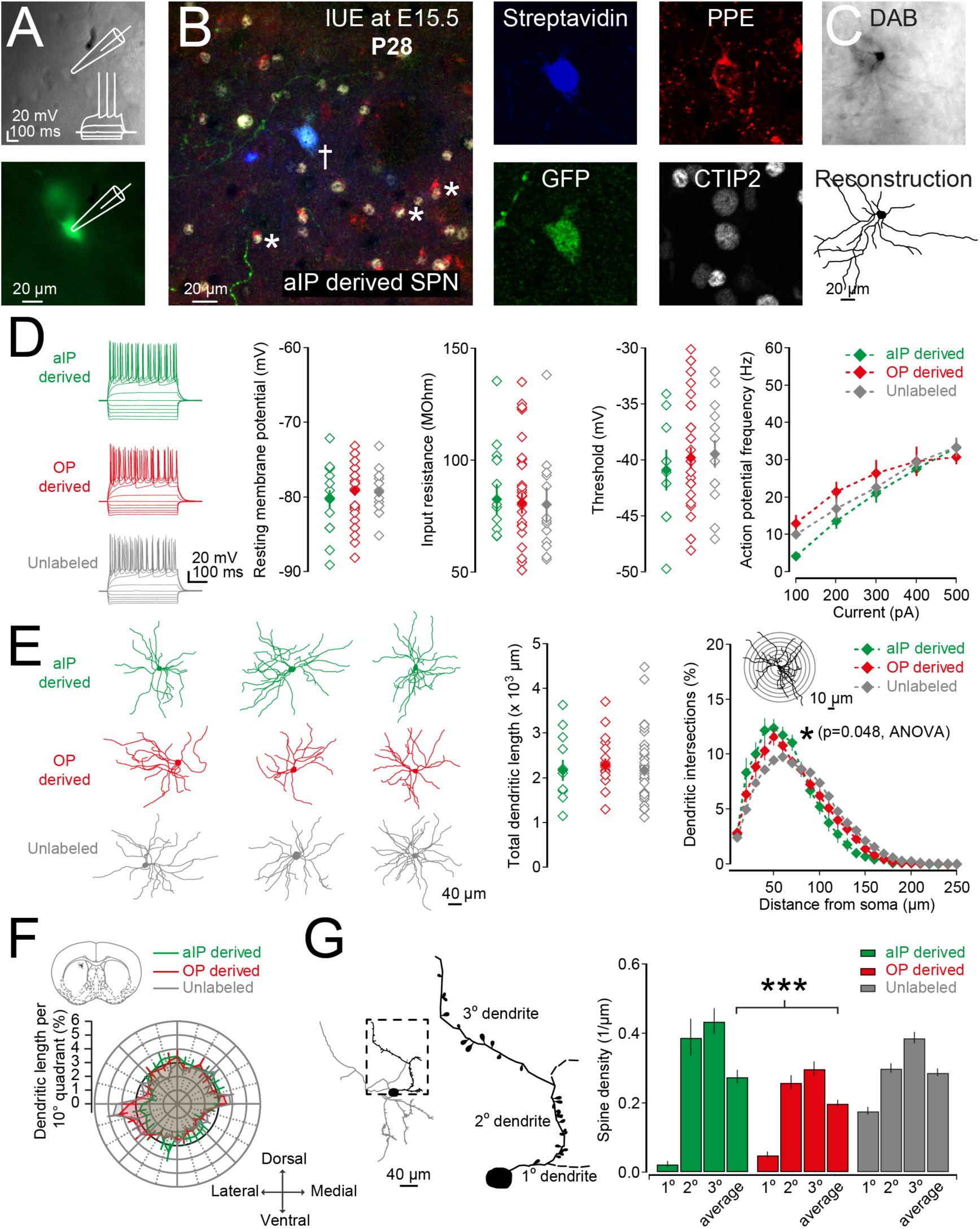
aIP derived SPNs exhibit greater dendritic complexity and spine density. (**A**) Electrophysiological properties of progenitor-derived striatal neurons were assessed using whole-cell patch-clamp recordings in acute brain slices. Example Dodt contrast image (top) and fluorescence image (bottom) of a recorded aIP derived GFP^+^ neuron. Inset: response of the neuron to hyperpolarizing and depolarizing current steps consistent with that of a SPN. (**B**) Recorded cells were labeled with biocytin during recordings and revealed using streptavidin-Alexafluor405 conjugated antibodies in fixed tissue (indicated with a cross) and tested for the expression of the SPN marker CTIP2 and the D2 SPN marker PPE. Indicated with white asterisks are neighboring CTIP2^+^ and PPE^+^ neurons. (**C**) Recorded cells were further processed for DAB immunohistochemistry allowing for reconstruction of their dendritic arbors as well as spine counts. (**D**) Hyperpolarizing and depolarizing current steps were used to characterise the electrophysiological properties of aIP derived, OP derived and unlabeled striatal neurons. The electrophysiological properties were consistent with those of SPNs and no significant differences were found between either group in their resting membrane potential, input resistance, spike threshold or action potential frequency. (**E**) Example reconstructions of DAB processed aIP derived (top), OP derived (middle) as well as unlabeled SPNs (bottom). The total dendritic length of the different SPNs did not differ. Assessment of dendritic complexity using Scholl analysis reveals that aIP derived neurons exhibit a subtle but significant larger dendritic complexity close to the soma. (**F**) Dendritic polarity analysis reveals that all SPNs exhibit a similar radial morphology. (**G**) Quantification of the number of spines on the primary (1°), secondary (2°) and tertiary (3°) dendrites reveals that aIP derived neurons exhibit an overall higher density of spines as compared to OP derived neurons.

**Table 1:**
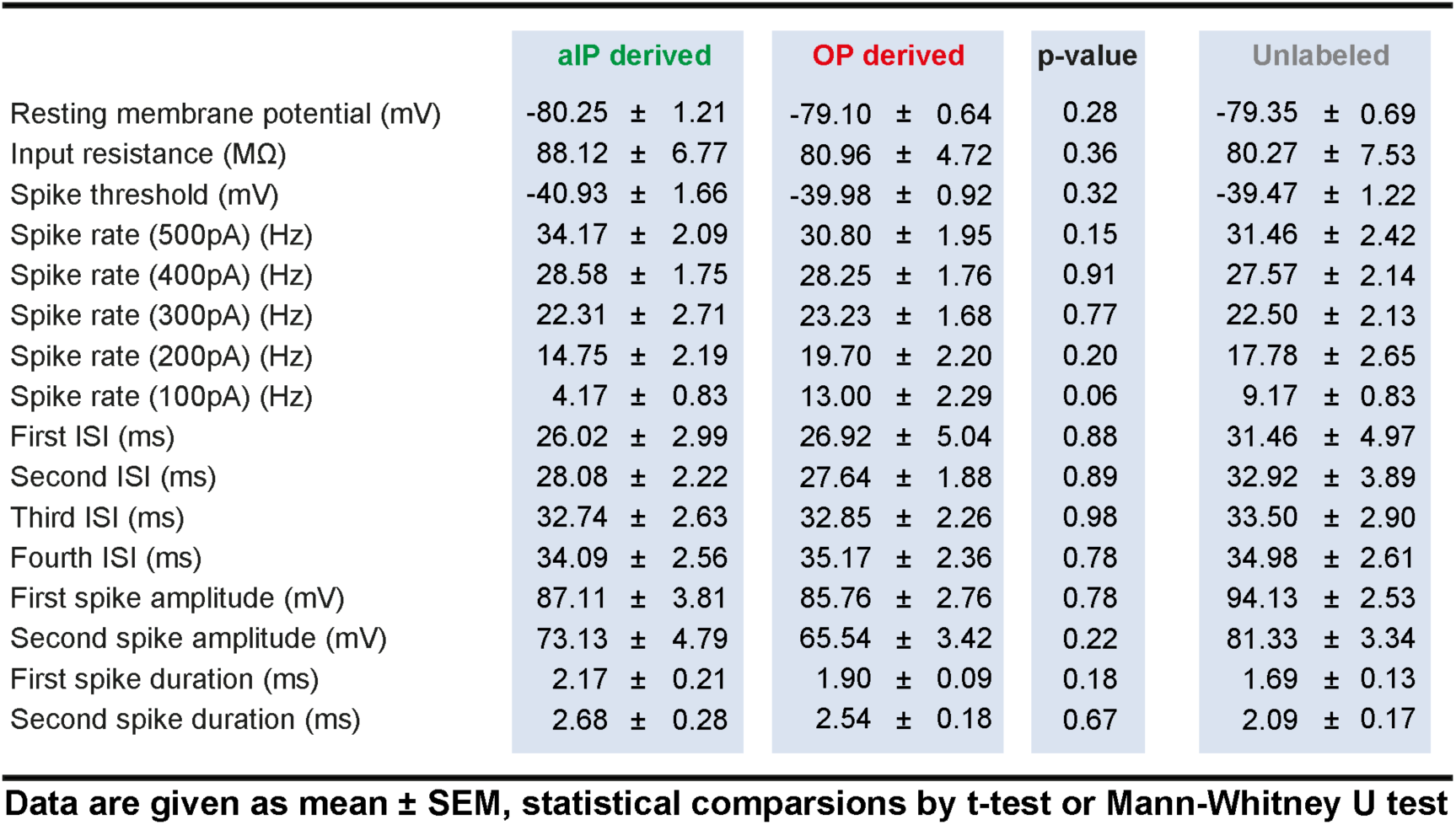
Intrinsic membrane properties of progenitor derived SPNs

|  | aIP derived | OP derived | p-value | Unlabeled |
| --- | --- | --- | --- | --- |
| Resting membrane potential (mV) | -80.25 ± 1.21 | -79.10 ± 0.64 | 0.28 | -79.35 ± 0.69 |
| Input resistance (MΩ) | 88.12 ± 6.77 | 80.96 ± 4.72 | 0.36 | 80.27 ± 7.53 |
| Spike threshold (mV) | -40.93 ± 1.66 | -39.98 ± 0.92 | 0.32 | -39.47 ± 1.22 |
| Spike rate (500pA) (Hz) | 34.17 ± 2.09 | 30.80 ± 1.95 | 0.15 | 31.46 ± 2.42 |
| Spike rate (400pA) (Hz) | 28.58 ± 1.75 | 28.25 ± 1.76 | 0.91 | 27.57 ± 2.14 |
| Spike rate (300pA) (Hz) | 22.31 ± 2.71 | 23.23 ± 1.68 | 0.77 | 22.50 ± 2.13 |
| Spike rate (200pA) (Hz) | 14.75 ± 2.19 | 19.70 ± 2.20 | 0.20 | 17.78 ± 2.65 |
| Spike rate (100pA) (Hz) | 4.17 ± 0.83 | 13.00 ± 2.29 | 0.06 | 9.17 ± 0.83 |
| First ISI (ms) | 26.02 ± 2.99 | 26.92 ± 5.04 | 0.88 | 31.46 ± 4.97 |
| Second ISI (ms) | 28.08 ± 2.22 | 27.64 ± 1.88 | 0.89 | 32.92 ± 3.89 |
| Third ISI (ms) | 32.74 ± 2.63 | 32.85 ± 2.26 | 0.98 | 33.50 ± 2.90 |
| Fourth ISI (ms) | 34.09 ± 2.56 | 35.17 ± 2.36 | 0.78 | 34.98 ± 2.61 |
| First spike amplitude (mV) | 87.11 ± 3.81 | 85.76 ± 2.76 | 0.78 | 94.13 ± 2.53 |
| Second spike amplitude (mV) | 73.13 ± 4.79 | 65.54 ± 3.42 | 0.22 | 81.33 ± 3.34 |
| First spike duration (ms) | 2.17 ± 0.21 | 1.90 ± 0.09 | 0.18 | 1.69 ± 0.13 |
| Second spike duration (ms) | 2.68 ± 0.28 | 2.54 ± 0.18 | 0.67 | 2.09 ± 0.17 |

**Data are given as mean ± SEM, statistical comparisons by t-test or Mann-Whitney U test**

We next investigated the morphological properties of the SPNs derived from aIP and OP progenitors. Recorded cells were post hoc processed for DAB immunohistochemistry allowing for reconstruction of their dendritic arbors as well as quantification of dendritic spines (Figure 3E - G). Comparing the total dendritic length we did not find significant differences in the length of their dendrites (aIP derived: 2184.7 ± 228.8 µm, OP derived: 2299.3 ± 147.2 µm and unlabeled neurons: 2182.6 ± 129.3 µm; p=0.626, Kruskal-Wallis H test, n = 11, 16 and 35 neurons, Figure 3E). We next investigated the dendritic complexity using Scholl analysis and find that all neurons exhibit the greatest dendritic complexity close to the soma (∼50 µm distance) with a small by significant greater complexity for aIP derived over OP derived neurons (p=0.048, ANOVA, n = 11 and 16, Figure 3E). Polarity analysis shows that the orientation of the dendrites is not significantly different between aIP derived, OP derived or unlabeled neurons and that all exhibit a radial morphology (Figure 3F). Quantification of dendritic spines on the primary, secondary and tertiary dendritic segments reveal that the aIP derived SPNs have a higher density of spines than the OP derived SPNs (average aIP derived: 0.28 ± 0.02 spines/µm and average OP derived: 0.20 ± 0.01 spines/µm, t-test, p=0.0058, n = 13 and 28, Figure 3G).

Combined these results suggest that both aIP and OP derived neurons exhibit all the electrical and morphological hallmarks of striatal SPNs. Furthermore, they show that subtle differences can be detected in the morphological properties of neurons derived from different progenitor pools (Guillamon-Vivancos *et al.*, 2019) in that aIP derived neurons exhibit greater local dendritic complexity and a higher density of dendritic spines.

### Integration of progenitor derived striatal SPNs into striatal neural circuits

We next asked to what extent aIP and OP derived neurons are integrated within the circuits of the basal ganglia and striatum. To do this we performed IUE at E15.5 to label both aIP derived and OP derived SPNs and examined both the synaptic inputs and outputs of labeled SPNs. We first investigated to what extent aIP and OP derived SPNs sample cortical afferents. Cortical afferents were activated using a stimulating electrode placed in the external capsule while recording from both aIP and OP derived neurons in the striatum (Figure 4B). All experiments were performed in the presence of GABA receptor antagonists to avoid erroneous activation of GABAergic afferents (see **Methods**). We find that aIP derived and OP derived SPNs exhibit similar amplitude EPSPs (aIP derived: 1.70 ± 0.44 mV and OP derived: 2.08 ± 0.62 mV, p=0.974, Mann Whitney test, n = 12 and 10, Figure 4D) and that these do not differ from amplitudes found in unlabeled SPNs (unlabeled: 1.50 ± 0.33 mV, aIP vs unlabeled: p=0.648 and OP vs unlabeled: p=0.650). However, we find that the duration and decay time of the EPSPs are significantly longer in the aIP derived neurons (Table 2) suggesting they might contain different configurations of glutamate receptors.

**Figure 4:**
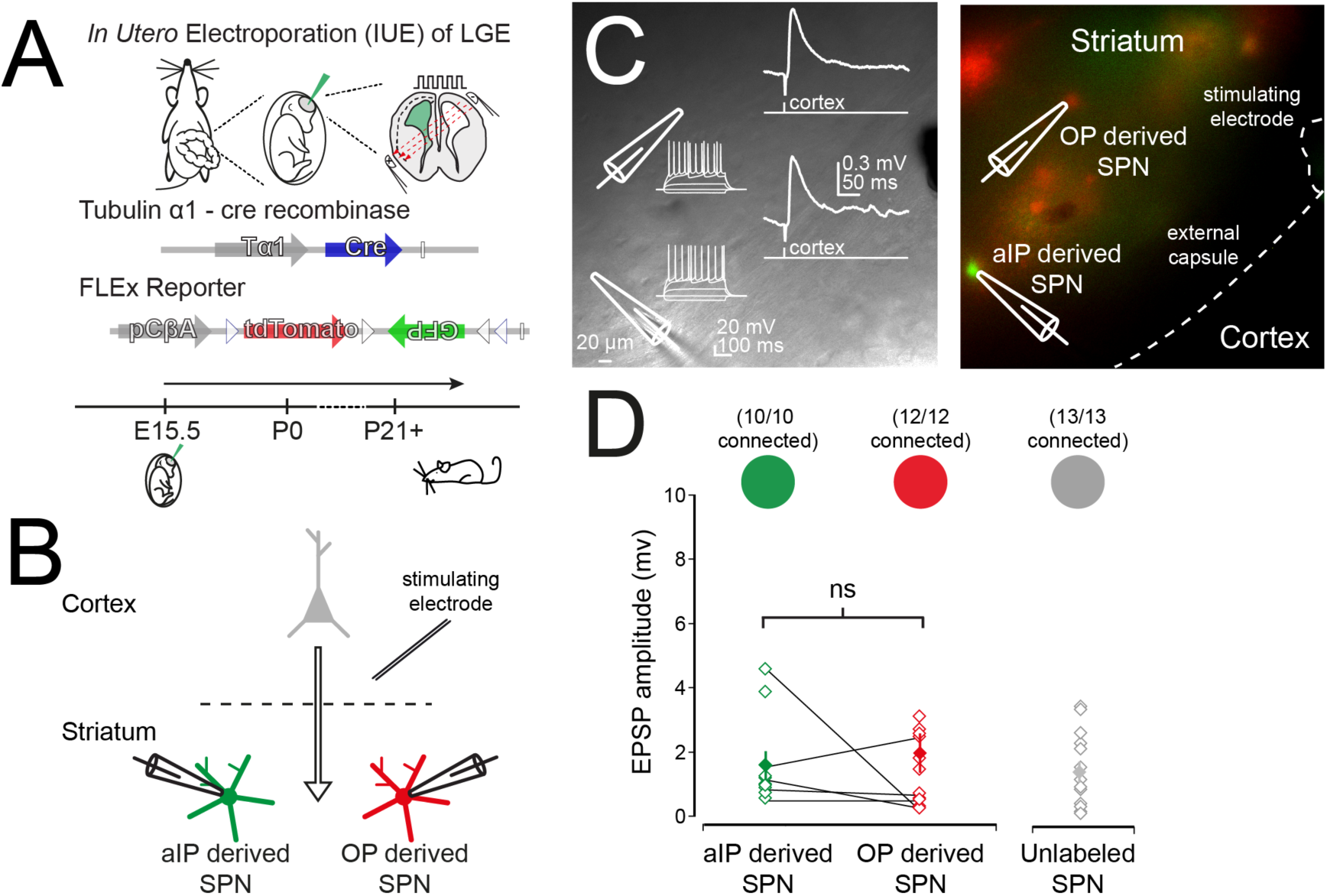
Both aIP and OP derived SPNs receive cortical input. (**A**) IUE of Tα1-Cre and FLEx reporter plasmids was performed at E15.5 to label both aIP and OP derived striatal neurons. (**B**) Diagram of the experimental setup consisting of electrical stimulation of cortical afferents while recording from both aIP and OP derived neurons in the striatum. All recordings were performed in the presence of GABA receptor antagonists. (**C**) Example Dodt contrast (left) and fluorescence (right) image of the experimental setup consisting of a Tungsten electrode placed in the external capsule and the simultaneous recording of a GFP^+^ aIP derived SPN and a TdTomato^+^ OP derived SPN. (**D**) Scatter plot of the amplitudes of the excitatory postsynaptic responses for aIP derived, OP derived and unlabeled SPNs upon stimulation of the cortical afferents. Note that all three classes of SPN receive similar amplitude cortically evoked EPSPs.

**Table 2:** Cortical synaptic response properties

|  | aIP derived | OP derived | p-value | Unlabeled |
| --- | --- | --- | --- | --- |
| Amplitude (pA) | 1.70 ± 0.44 | 2.08 ± 0.62 | 0.77 | 1.50 ± 0.33 |
| Duration (ms) | 143.9 ± 14.48 | 99.30 ± 10.47 | 0.03 | 112.81 ± 11.76 |
| Rise time (ms) | 5.73 ± 0.42 | 5.87 ± 0.95 | 0.83 | 4.47 ± 0.29 |
| Decay time (ms) | 66.20 ± 6.98 | 44.63 ± 5.32 | 0.03 | 55.86 ± 5.90 |

**Data are given as mean ± SEM, statistical comparisons by Mann-Whitney test.**

We next asked to what extent these progenitor derived neurons innervate neighbouring SPNs and in particular whether aIP derived neurons similarly innervate the D1-expressing direct pathway and D2-expressing indirect pathway SPNs. We used electroporation of Tα1-cre and creON-mCherry plasmids (Saunders *et al.*, 2012) at E15.5 to selectively express the light-activatable channel ChR2 in aIP derived neurons in C57Bl/6 and D1 or D2-GFP transgenic mice with the latter facilitating the classification of patched neurons as either D1 or D2 SPNs (Figure 5A).

**Figure 5:**
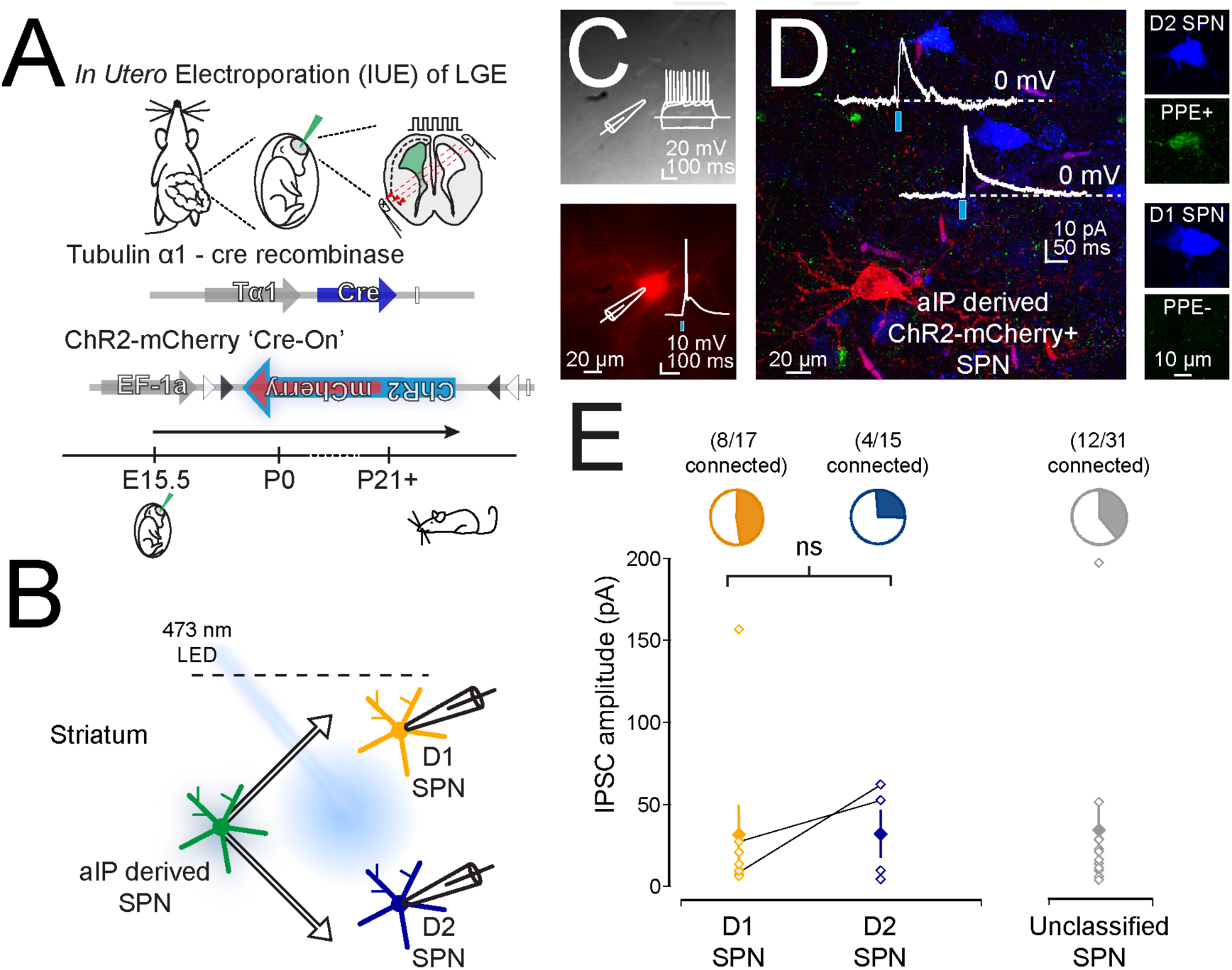
aIP derived SPNs connect to both D1 and D2 SPNs. (**A**) IUE of Tα1-Cre and DIO-ChR2-mCherry plasmids allowed for the expression of ChR2-mCherry in aIP derived neurons. (**B**) Diagram of the experimental setup consisting of recordings of D1 and D2 SPNs while stimulating aIP derived neurons with a 473 nm wide-field LED. All recordings were performed in the presence of glutamate receptor antagonists. (**C**) Whole-cell patch-clamp recordings of aIP derived ChR2-mCherry expressing striatal neurons revealed that ∼3 ms flashes of light could elicit single action potentials. Inset top: response of the neuron to hyperpolarising and depolarising current steps. (**D**) To investigate striatal inhibitory synaptic output from aIP derived neurons, simultaneous whole-cell patch-clamp recordings were performed from D1 and D2 SPNs, as determined during recordings in D1 or D2-GFP transgenic mice or posthoc using immunocytochemistry for PPE. Example image of streptavidin labeled PPE^+^ (upper neuron) and PPE^-^ SPN (lower neuron) in close apposition to an aIP derived-ChR2-mCherry^+^ SPN. Traces correspond to light evoked IPSCs. (**E**) aIP derived striatal neurons exhibited a slight preference to connect to D1 SPNs (47%) and optical activation of aIP derived neurons results in similar amplitude IPSCs in all tested D1, D2 and unclassified SPNs.

In postnatal acute brain slices we were now able to deliver short pulses of blue light to activate ChR2-mCherry expressing aIP derived neurons (Figure 5C) while performing whole-cell voltage-clamp recordings of single or pairs of D1 and D2 SPNs at a holding potential of 0 mV to facilitate detection of inhibitory events (Figure 5B and D). We recorded from a total of 63 SPNs of which 17 were confirmed D1 SPNs, 15 were confirmed D2 SPNs and 31 were unclassified. Although we find that the incidence of finding a connected SPN is slightly higher for the D1 SPNs than the D2 SPNs (D1 SPN: 47.1% and D2 SPN: 26.7%) this does not reach statistical significance (p=0.517; Fisher’s exact test). For the neurons that received inputs from the aIP derived neurons we find no significant difference in the amplitude of the light evoked inhibitory postsynaptic current (IPSC) as recorded at a holding potential of 0 mV (D1: 31.23 ± 18.11 pA, D2: 31.97 ± 14.69 pA and unclassified: 34.15 ± 15.30 pA, p=0.797, Kruskall Wallis test, n = 8, 4 and 12, Figure 5E).

In conclusion, these results suggest that both aIP and OP derived SPNs are embedded within the neural circuits of the striatum. Indeed, both aIP and OP derived striatal SPNs receive excitatory glutamatergic cortical input and aIP derived neurons send inhibitory GABAergic outputs to neighbouring D1 and D2 SPNs. Interestingly, we find that aIP and OP derived neurons differentially respond to cortical inputs, with aIP derived SPNs exhibiting prolonged EPSP kinetics, suggesting that they might sample excitatory inputs differently.

## Discussion

In this study we examined to what extent neural progenitor pool diversity in the lateral ganglionic eminence (LGE) contributes to cellular and circuit diversity in the striatum. Using *in utero* electroporation we are able to label two distinct pools of dividing neural progenitors in the LGE, which are active during later stages of neurogenesis. Using differential expression of the tubulin alpha1 (Tα1) promoter we are able to label a Tα1-expressing pool of neural progenitors we refer to as apical intermediate progenitors (aIPs) and a non-Tα1-expressing pool we refer to as other progenitors (OPs). We find that both progenitor types are actively dividing in the ventricular zone of the LGE around embryonic days (E)15.5-16.5, but exhibit distinct proliferative behaviours. Whereas aIP progenitors exhibit a short round morphology during division, lack a basal process and exhibit fast cell cycle kinetics, OP progenitors tend to retain a basal process and exhibit slower cell cycle kinetics. We show that both can generate D1-expressing direct pathway spiny projection neurons (SPNs) as well as D2-expressing indirect pathway SPNs found in both patch and matrix compartments of the striatum. Whereas aIP and OP derived SPNs were found intermingled throughout the striatum on average the aIP derived population of SPNs inhabited more medial aspects of the striatum. Whole-cell patch-clamp recordings and morphological reconstruction of aIP and OP derived neurons confirmed they were SPNs with aIP derived SPNs exhibiting a more complex local dendritic tree and an increased dendritic spine density. Electrical circuit mapping shows that both aIP and OP derived SPNs sample input from cortical excitatory afferents with aIP derived SPNs exhibiting differential sampling and longer duration responses. Lastly, optogenetic circuit mapping reveals that aIP derived SPNs send local GABAergic outputs to both D1 and D2 SPNs. Overall, we show that it is possible to label diverse pools of neural progenitors in the LGE and their progeny in the striatum. Diversity in the pool of progenitors can generate diversity in adult striatal neurons, which manifests itself in different spatial distributions, morphological properties and sampling of excitatory afferents.

### Different pools of embryonic neural progenitors in the LGE

The neural progenitors in the LGE of the embryonic mouse brain (Smart, 1976) generate most of the SPNs of the striatum as well as some of the interneurons of the olfactory bulb (Olsson *et al.*, 1998; Stenman *et al.*, 2003) with postnatal generation of striatal neurons also contributing (Stopczynski *et al.*, 2008). Recent studies have highlighted that the LGE contains as much heterogeneity in neural progenitors (Pilz *et al.*, 2013) as pallial structures (Franco & Muller, 2013). Indeed, in the LGE these progenitors include radial glial cells, basal radial glial cells, basal progenitors, subapical progenitors as well as short neural precursors, amongst others (Pilz *et al.*, 2013). It is currently unknown how this heterogeneity at the progenitor level relates to the cellular and circuit diversity found in the striatum.

Several lines of evidence suggest that that the population of progenitors labeled using the promoter sequence for Ta1 correspond to the short neural precursors and subapical progenitors recently described in the LGE (Gal *et al.*, 2006; Stancik *et al.*, 2010; Pilz *et al.*, 2013; Ellender *et al.*, 2018). We find that the population of Tα1-expressing neural progenitors divides at some distance from the ventricular wall but mostly with the ventricular zone, their cell cycle kinetics are fast and correspond to about 17 hours, they form a large population of dividing progenitors during later stages of neurogenesis (E15.5-E16.5) and lastly they lack a basal process during division. These criteria are consistent to those described for short neural precursors and the subapical progenitors that are often generated from the short neural precursors (Pilz *et al.*, 2013). Conversely, even though we find that our population of non-Tα1-expressing neural progenitors also divides in the ventricular zone, it is more dominant during early stages of neurogenesis (E12.5/E13.5), has slower cell cycle kinetics corresponding to about 20 hours (Pilz *et al.*, 2013), and many progenitors retain a basal process during division. These criteria are consistent with the described morphology of radial glial cells in the LGE (Pilz *et al.*, 2013) and the cortex (Gal *et al.*, 2006; Kriegstein & Alvarez-Buylla, 2009).

As the Tα1-expressing neural progenitors could be found actively dividing in the apical aspects of the ventricular zone and had many properties of both short neural precursors and subapical progenitors (Pilz *et al.*, 2013) we proposed to refer to them collectively as apical intermediate progenitors (aIPs) and our non-Tα1-expressing neural progenitors as other progenitors (OPs). Therefore, at the point of labeling, our *in utero* strategy marked a progenitor population enriched for aIPs and a population of concurrently dividing OPs. Although the method allowed us to distinguish neurons derived from different progenitor pools, it provides limited information on the lineage pathways taken by the neurons. Moreover, neural progenitor lineage in the LGE is complex with extended proliferative divisions and likely both progenitor pools generate neurons via other progenitors (Pilz *et al.*, 2013).

### Progenitor derived striatal spiny projection neurons and associated circuits

We find that both aIP and OP neural progenitors generate neurons that can be found intermingled in the striatum. By varying the embryonic time at which we perform our IUE we confirm that aIP progenitors are mainly active at later stages of neurogenesis as we find more aIP derived neurons when we electroporate at E15.5 as compared to E12.5. Immunohistochemistry for the SPN marker CTIP2 (Arlotta *et al.*, 2008) reveals that most if not all labeled neurons are SPNs independent of whether IUE was performed at E12.5 or E15.5. Furthermore, immunohistochemistry for the D2 SPN marker PPE (Gerfen *et al.*, 1990) reveals that both aIP and OP progenitor pools can generate D1 and D2 SPNs at both E12.5 and E15.5. These observations are consistent with the idea that the generation of D1 and D2 SPNs is controlled by other genetic factors, environmental factors or molecular clocks (Lobo *et al.*, 2006; Franco *et al.*, 2012; Cepko, 2014; Kelly *et al.*, 2018). Indeed, similar observations have been made in the cortex where both layer 4 stellate neurons and layer 2/3 pyramidal neurons can be generated from Tα1-expressing as well as GLAST-expressing neural progenitors (Stancik *et al.*, 2010) and indeed a single cortical progenitor can generate clones with layer-specific characteristics in all layers of cortex (Yu *et al.*, 2009). Whereas the neural progenitor pool of origin did not seem to affect SPN type or location in patch or matrix compartments (Pert *et al.*, 1976; Gerfen, 1984; Herkenham *et al.*, 1984; Gerfen *et al.*, 1985; Jimenez-Castellanos & Graybiel, 1989; Gerfen, 1992; Crittenden & Graybiel, 2011), we find that the earlier time of electroporation resulted in both pools of progenitors generating a higher proportion of D1 SPNs (Marchand & Lajoie, 1986; van der Kooy & Fishell, 1987; Kelly *et al.*, 2018) as well as generating more SPNs in patch compartments (Brand & Rakic, 1979; Graybiel & Hickey, 1982; van der Kooy & Fishell, 1987; Fishell *et al.*, 1990; Newman *et al.*, 2015; Tinterri *et al.*, 2018).

Interestingly, we find that the different pools of neural progenitors generate D1 and D2 SPNs found in different aspects of the striatum, with aIP derived SPNs on average found in more medial aspects of the striatum. Already at embryonic ages the aIPs and related young neurons are found at slightly more distal locations relative to the ventricle, which might suggest that the location of aIP derived neurons in the adult brain results from such early differences. However, migration in the striatum is complex in that newly born neurons undergo tangential, radial and other forms of migration (Halliday & Cepko, 1992; Tan & Breen, 1993; Reid & Walsh, 2002; Tinterri *et al.*, 2018), suggesting other factors could also contribute to the ultimate location of striatal neurons (Franco *et al.*, 2012; Kelly *et al.*, 2018). Interestingly, detailed studies of the inputs to the striatum have revealed that dorsomedial aspects of the striatum exhibit a high degree of input heterogeneity (Pan *et al.*, 2010; Guo *et al.*, 2015; Hunnicutt *et al.*, 2016) suggesting these different progenitor derived SPNs might process distinct inputs.

Electrophysiological and morphological study of single neurons reveals that both aIP and OP derived neurons mostly exhibit similar properties consistent with those of SPNs (Day *et al.*, 2008; Gertler *et al.*, 2008; Krajeski *et al.*, 2018). Interestingly, we find that aIP derived neurons exhibit a greater local dendritic complexity and a higher dendritic spine density than OP derived neurons. This difference might suggest that they differentially sample excitatory inputs. Indeed, whereas the overall strength of cortical inputs to both aIP and OP derived neurons seems similar, the aIP derived neurons exhibit comparatively long duration EPSPs. This might well result from the expression of a different complement of glutamate receptors postsynaptically.

Lastly, it is known that SPNs in the striatum form strong lateral inhibitory connections with each other with which they can regulate each other’s activity (Taverna *et al.*, 2008; Planert *et al.*, 2010; Chuhma *et al.*, 2011; Krajeski *et al.*, 2018) and these are not uniform but depend on the type of pre- and postsynaptic SPN (Taverna *et al.*, 2008; Planert *et al.*, 2010; Krajeski *et al.*, 2018). For example, the D2 SPNs form a large number of strong reciprocal connections with each other (Planert *et al.*, 2010; Krajeski *et al.*, 2018). Although our current results suggest that aIP derived SPNs innervate both local D1 and D2 SPNs, future work will further inform us on whether and how diverse neural progenitor pools contribute to the establishment of further fine-scale neural circuits in the striatum.

In conclusion, we show that we are able to fluorescently pulse-label two different pools of neural progenitors in the LGE and their offspring. Using this method we reveal that heterogeneity in neural progenitor pools in the LGE can contribute to diversity in the spiny projection neurons and circuits of the striatum. In particular, we find that aspects of spatial location and morphology of neurons, as well as their sampling of excitatory afferents seem particularly sensitive to neural progenitor origin. Future investigations will be informative as to what extent they contribute to other aspects of striatal diversity (Grillner & Robertson, 2016) and whether neural progenitor dysregulation plays a role in early onset neurodevelopmental disorders such as OCD, autism and Tourette’s syndrome (Graybiel & Rauch, 2000; Del Campo *et al.*, 2011; Langen *et al.*, 2011; McNaught & Mink, 2011; Shepherd, 2013; Albin, 2018).

## Acknowledgements

We would like to thank members of the Ellender lab for advice and comments. We gratefully acknowledge Ulrich Müller and Tarik Haydar for providing reagents, Peter Somogyi, Peter Magill and Colin Akerman for providing access to equipment, Ben Micklem for providing technical assistance and Monzilur Rahman, Rebecca Waterfield, Nicholas Pasternack and Eoin Mac Reamoinn for initial help. TJE was supported by a MRC Career Development Award (MR/M009599/1) and AMD by an Imperial College research bursary.

## Methods & Materials

### Animals

All experiments were carried out on C57/BL6 wildtype and heterozygous D1-GFP or D2-GFP mice of both sexes with *ad libitum* access to food and water. The D1-GFP or D2-GFP BAC transgenic mice report subtypes of the dopamine receptor, either D1 or D2, by the presence of GFP (Mutant Mouse Regional Resource Centers, MMRRC). Details of the mice and the methods of BAC mice production have been published (Gong *et al.*, 2003) and can be found on the GENSAT website [GENSAT (2009) The Gene Expression Nervous System Atlas (GENSAT) Project. In: NINDS, Contracts N01NS02331 and HHSN271200723701C, The Rockefeller University (New York), http://www.gensat.org/index.html]. The BAC transgenic mice were backcrossed to a C57/BL6 background over 20+ generations prior to use and kept as a heterozygous mouse line to avoid published issues using these transgenic lines (Bagetta *et al.*, 2011; Kramer *et al.*, 2011; Chan *et al.*, 2012; Nelson *et al.*, 2012). All mice were bred, IVC housed in a temperature controlled animal facility (normal 12:12 h light/dark cycles) and used in accordance with the UK Animals (Scientific Procedures) Act (1986). Females were checked for plugs daily; the day of the plug was considered embryonic day (E)0.5 and injection of plasmids and *in utero* electroporation (IUE) was performed at E12.5 - E16.5.

### Electroporation

*In utero* electroporation (IUE) was performed using standard procedures. In short, pregnant females were anaesthetized using isoflurane and their uterine horns were exposed by midline laparotomy. A mixture of plasmid DNA (∼1.5 µg/µl) and 0.03% fast green dye was injected intraventricularly using pulled micropipettes through the uterine wall and amniotic sac. Plasmid DNA included: (i) ‘Tα1-Cre’, in which the gene for Cre recombinase is under the control of a portion of the Tα1 promoter (Stancik *et al.*, 2010); (ii) ‘CβA-FLEx’ which uses the chicken β-*actin* promoter to control a flexible excision (FLEx) cassette in which Cre recombination switches expression from TdTomato fluorescent protein to enhanced green fluorescent protein (Franco *et al.*, 2012); and (iii) ‘DIO-ChR2-mCherry’ (pAAV-EF1a-doublefloxed-hChR2(H134R)-mCherry-WPRE-HGHpA; Addgene #20297), in which Cre recombination turns on the expression of channelrhodopsin-2 (ChR2) under the control of the human elongation factor-1a promoter (Saunders *et al.*, 2012). Total volume injected per pup was ∼1 µl. Tα1-Cre and CβA-FLEx constructs (and other combinations of constructs) were injected as a 1:1 ratio of plasmid DNA (each 3 µg/µl, so final concentration of both constructs was 1.5 µg/µl). The cathode of the Tweezertrode (Genetronics) was placed just above the eye outside the uterine muscle and the anode was placed slightly lower at the contralateral cheek region (Baumgart & Baumgart, 2016). Five pulses (50 ms duration separated by 200 - 950 ms) at 42-60V with a BTX ECM 830 pulse generator (Genetronics). Typically around 80% of the pups underwent electroporation. Afterwards the uterine horns were placed back inside the abdomen, the cavity filled with warm physiological saline and the abdominal muscle and skin incisions were closed with vicryl and prolene sutures, respectively. Dams were placed back in a clean cage and monitored closely until the birth of the pups.

### Slice preparation and recording conditions

Acute striatal slices were made from postnatal animals at 3-5 weeks of age. Animals were anaesthetized with isoflurane and then decapitated. Coronal 350-400 µm slices were cut using a vibrating microtome (Microm HM650V). Slices were prepared in artificial cerebrospinal fluid (aCSF) containing (in mM): 65 Sucrose, 85 NaCl, 2.5 KCl, 1.25 NaH_2_PO_4_, 7 MgCl_2_, 0.5 CaCl_2_, 25 NaHCO_3_ and 10 glucose, pH 7.2-7.4, bubbled with carbogen gas (95% O_2_ / 5% CO_2_). Slices were immediately transferred to a storage chamber containing aCSF (in mM): 130 NaCl, 3.5 KCl, 1.2 NaH_2_PO_4_, 2 MgCl_2_, 2 CaCl_2_, 24 NaHCO_3_ and 10 glucose, pH 7.2 - 7.4, at 32 °C and bubbled with carbogen gas until used for recording. Striatal slices were transferred to a recording chamber and continuously superfused with aCSF bubbled with carbogen gas with the same composition as the storage solution (32 °C and perfusion speed of 2 ml/min). Whole-cell recordings were performed using glass pipettes (∼6MΩ), pulled from standard wall borosilicate glass capillaries and containing for whole-cell current-clamp (in mM): 110 potassium gluconate, 40 HEPES, 2 ATP-Mg, 0.3 Na-GTP, 4 NaCl and 4 mg/ml biocytin (pH 7.2-7.3; osmolarity, 290-300 mosmol/l) and for whole-cell voltage-clamp (in mM): 120 cesium gluconate, 40 HEPES, 4 NaCl, 2 ATP-Mg, 0.3 Na-GTP, 0.2 QX-314 and 4 mg/ml biocytin (pH 7.2–7.3; osmolarity, 290-300 mosmol/L). Recordings were made using Multiclamp 700B amplifiers and filtered at 4kHz and acquired using an InstruTECH ITC-18 analog/digital board and WinWCP software (University of Strathclyde, RRID:SCR_014713) at 10 kHz.

### Stimulation and recording protocols

Hyperpolarizing and depolarizing current steps were used to assess the intrinsic properties of the recorded SPNs including input resistance and spike threshold (using small incremental current steps) as well as the properties of action potentials (amplitude, frequency and duration). Properties were assessed immediately on break-in. Activation of excitatory cortical afferents was performed via a bipolar stimulating electrode (FHC Inc., USA) placed in the external capsule, and in the presence of blockers of inhibitory GABAergic transmission including the GABA_A_-receptor antagonist SR95531 (1 µM) and the GABA_B_-receptor antagonist CGP52432 (2 µM). Afferents were activated every 5s with up to 20 repetitions and excitatory postsynaptic potentials (EPSPs) were recorded from the patched SPNs. Photoactivation of ChR2 was achieved using widefield 2-3 ms duration light pulses of ∼1 mW via a TTL triggered CoolLED pE-300 system (CoolLED, Andover, UK)

### Analysis of recordings

Data were analyzed offline using custom written programs in Igor Pro (Wavemetrics, RRID:SCR_000325). The input resistance was calculated from the observed membrane potential change after hyperpolarizing the membrane potential with a set current injection. The spike threshold was the membrane voltage at which the SPN generated an action potential. The action potential amplitude was taken from the peak amplitude of the individual action potentials relative to the average steady-state membrane depolarization during positive current injection. Action potential duration was taken as the duration between the upward and downward stroke of the action potential at 25% of the peak amplitude. Evoked EPSPs and IPSCs were defined as upward or downward deflections of more than 2 standard deviations (SD) on average synaptic responses generated after filtering and averaging original traces (0.1 Hz high-pass filter and 500 Hz low-pass filter) and used for analysis of synaptic properties. Synaptic properties include measurements of peak amplitude, duration (measured from the start of the upward/downward stroke of the event until its return to the pre-event baseline), rise time (time between 20% and 80% of the peak amplitude) and decay time (measured as the time from peak amplitude until the event returned to 50% of peak amplitude).

### Histological analyses

Following whole-cell patch-clamp recordings the brain slices were fixed in 4% paraformaldehyde in 0.1 M phosphate buffer (PB; pH 7.4). Biocytin-filled neurons were visualized by incubating sections in 1:10,000 streptavidin AlexaFluor405-conjugated antibodies (ThermoFisher Scientific, Cat#:S32351). Visualized neurons were labeled for chicken ovalbumin upstream promoter transcription-factor interacting protein-2 (CTIP2, 1:1000, rat, Abcam, Cat#:ab14865, RRID:AB_2064130) and pre-proenkephalin (PPE, 1:1000, rabbit, LifeSpan Biosciences, Cat#:LS-C23084, RRID:AB_902714) in PBS containing 0.3% Triton X-100 (PBS-Tx) overnight at 4^◦^C followed by incubation with goat-anti-rat AlexaFluor647 (1:500; ThermoFisher Scientific, CAT#:A-21247, RRID:AB_141778) and goat-anti-rabbit AlexaFluor555 (1:500; ThermoFisher Scientific, CAT#:A-21429, RRID:AB_2535850) or goat-anti-rabbit AlexaFluor488 (1:500; ThermoFisher Scientific, CAT#:A32731, RRID:AB_2633280) secondary antibodies in 0.3% PBS-Tx for 2 h at RT for D1 or D2 SPN classification. Occasionally the endogenous fluorescence would be boosted with antibodies against GFP (1:1000, chicken, Aves Labs, CAT# GFP-1020, RRID:AB_10000240) or TdTomato (1:1000; rat; anti-RFP; Chromotek, CAT# 5f8-100, RRID:AB_2336064) or slices were co-stained with the nuclear marker 4′,6-diamidino-2-phenylindole (DAPI) in PBS (1:100,000) to facilitate the delineation of brain structures. CTIP2 is expressed by SPNs and not interneurons (Arlotta *et al.*, 2008) and PPE reliably labels indirect pathway D2 SPNs (Lee *et al.*, 1997; Sharott *et al.*, 2017). PPE antibody staining was facilitated through antigen retrieval by heating sections at 80^◦^ C in 10 mM sodium citrate (pH 6.0) for approximately 30-60 min prior to incubation with PPE primary antibody. After classification of SPNs the slices were washed 3 times in PBS and processed for DAB immunohistochemistry using standard procedures.

Whole-brain fixation of embryonic and adult brains was performed by rapid decapitation of the head and submersion in oxygenated sucrose cutting solution before submersion in 4% paraformaldehyde in 0.1 M phosphate buffer (PB; pH 7.4). The brains were fixed for 24 – 48 hours, after which they were washed in PBS. Whole-brain tissue was directly, or in the case of embryonic tissue after embedding in 5% agar, sectioned at 50 µm on a vibrating microtome (VT1000S; Leica Microsystems). All sections were pre-incubated in 10-20% normal donkey serum (NDS; Vector Laboratories) or normal goat serum (NGS; Vector Laboratories) in PBS for more than 1h at RT. GFP^+^ (Tα1^+^) and TdTomato^+^ (Tα1^-^) progenitors and neurons were often visualized without antibody-mediated augmentation of fluorescence but in rare cases the endogenous fluorescence was boosted with antibodies against GFP (1:1000, chicken, Aves Labs, CAT#:GFP-1020, RRID:AB_10000240) or TdTomato (1:1000; rat; anti-RFP; Chromotek, CAT#:5f8-100, RRID:AB_2336064) and Goat-Anti-chicken AlexaFluor488 (1:500; Life Technologies, CAT#:A11039, RRID:AB_142924) and Goat-Anti-rat AlexaFluor555 (1:500; Life Technologies, CAT#:A-21429, RRID:AB_2535850). In embryonic tissue antibody labeling was used to label pH3 in neural progenitors (1:500; rabbit; Millipore, CAT#:06-570, RRID:AB_310177). Adult tissue was either co-stained in 1:100,000 DAPI in PBS to facilitate the delineation of brain structures or was labeled for MOR (1:3000, goat, ImmunoStar CAT#:24216, RRID:AB_572251) or CTIP2 (1:1000, rat: Abcam Cat#:ab14865, RRID:AB_2064130) and PPE (1:1000, rabbit, LifeSpan Biosciences Cat#:LS-C23084, RRID:AB_902714). PPE staining was facilitated through antigen retrieval by heating sections at 80^◦^ C in 10 mM sodium citrate (pH 6.0) for approximately 30 min prior to incubation with 1:1000 rabbit anti-PPE in PBS-Tx and 1% NDS overnight at 4^◦^C, after which the reaction was revealed by incubating with 1:500 donkey-anti-rabbit AlexaFluor647 (1:500, Life Technologies CAT#:A31573, RRID:AB_2536183) in PBS-Tx for 2 h at RT.

### Stereology and analysis of tissue

Fluorescence images were captured with a LSM 710 confocal microscope using ZEN software (Zeiss, RRID:SCR_013672) or Leica DM5000B epifluorescence microscope using Openlab software (PerkinElmer, RRID:SCR_012158). Counting of labeled GFP^+^ and TdTomato^+^ progenitors and young neurons and assessing their location within the embryonic brain was performed using ImageJ (RRID:SCR_003070) on z-stacks of ∼40 µm thickness. In embryonic tissue occasionally yellow cells could be seen which were counted as GFP^+^ and were assumed to have undergone recombination relatively recently. Positive cells had a fluorescence signal that was at least twice the background fluorescence (measured from randomly selected regions of the tissue). X- and y-coordinates labeled cells were used to calculate the distance from the ventricle and spread. Counting of progenitor cell basal processes was performed in z-stack projections of confocal stacks of ∼40 µm thickness. All clearly delineated processes above the SVZ and extending towards the pial surface were counted. M-phase reentry after IUE for aIP and OP was estimated from co-labeling of cells with the mitotic marker pH3 in tissue fixed with varying time-delays after IUE of Tα1-Cre and CβA-FLEx plasmids (Stancik *et al.*, 2010). Localizing GFP^+^ and TdTomato^+^ progenitors and young neurons in various sub regions of the LGE was performed using a combination of anatomical landmarks (Schambra & Schambra, 2008) and previous delineations (Flames *et al.*, 2007). Olfactory bulb analysis was performed using a total of 5 brains and all GFP or TdTomato positive cells were counted in z-stacks of ∼40 µm thickness.

Progenitor derived neuron counting and analysis was performed similar to (Garas *et al.*, 2016; Garas *et al.*, 2018). In brief, a version of design-based stereology, the ‘modified optical fractionator’ was used to generate unbiased cell counts and map distributions of striatal neurons in rostral, middle and caudal sections (Franklin & Paxinos, 2008). Once the chosen striatal coronal planes were identified and the immunofluorescence protocol carried out, the dorsal striatum was delineated using a Zeiss Imager M2 epifluorescence microscope (Carl Zeiss, AxioImager.M2) equipped with a 20X (Numerical Aperture = 0.8) objective and StereoInvestigator v9.0 software (MBF Biosciences). Imaging was subsequently performed by capturing a series of completely tessellated, z-stacked images (each 1 µm thick) at depths from 2 to 12 µm from the upper surface of each section at the level of the striatum (thereby defining a 10 µm-thick optical disector). As counts were performed across the entirety of the dorsal striatum within a given rostro-caudal plane, the grid size and counting frame were set to the same size of 420 x 320 mm. To minimize confounds arising from surface irregularities, neuropil within a 2 µm ‘guard zone’ at the upper surface was not imaged. A neuron was counted if the top of its nucleus came into focus within the disector. If the nucleus was already in focus at the top of the 10 µm -thick optical disector the neuron was excluded. Normalised positions were calculated as described in (Garas *et al.*, 2016; Garas *et al.*, 2018). Mediolateral and dorsoventral bias within each individual section was assessed by computing a Wilcoxon Sign rank test on the positions of all neurons across or within groups to test whether they significantly differed from zero (minimum 8 neurons for a given section). Mediolateral and dorsoventral positions of red and green neurons across animals were compared by computing a Wilcoxon sign rank test on the normalised position in each direction for each section, when there were a minimum of 8 neurons of each type in a single section (n = 20 sections).

DAB-immunoreactive neurons were visualized on a brightfield microscope and were reconstructed and analyzed using Neurolucida and Neuroexplorer software (MBF Bioscience, RRID:SCR_001775). Only labeled neurons that exhibited a full dendritic arbor were included for analysis e.g. cells with clear truncations were not included in the dataset. Scholl analysis and polarity analysis was performed using standard procedures. In brief, both Scholl and polarity plots were generated for individual SPNs by calculating the total dendritic length located within 10° segments with increasing distance from the soma. The dendritic lengths were subsequently normalised for an individual SPN and averaging the normalised plots of individual neurons generated final plots.

### Statistics

All data are presented as means ± SEM. The ‘n’ refers to the number of brains (Figure 1 and 2) or neurons (Figure 3 – 5) tested. Statistical tests were all two-tailed and performed using SPSS 17.0 (IBM SPSS statistics, RRID:SCR_002865) or GraphPad Prism version 5.0 (GraphPad software, RRID:SCR_002798). Synaptic connectivity ratios were compared with Fisher’s Exact test. Continuous data were assessed for normality and appropriate parametric (ANOVA, paired t-test and unpaired t-test) or non-parametric (Mann-Whitney U) statistical tests were applied (* p<0.05, ** p<0.01, *** p<0.001).

---

**Supplementary Figure 1:**
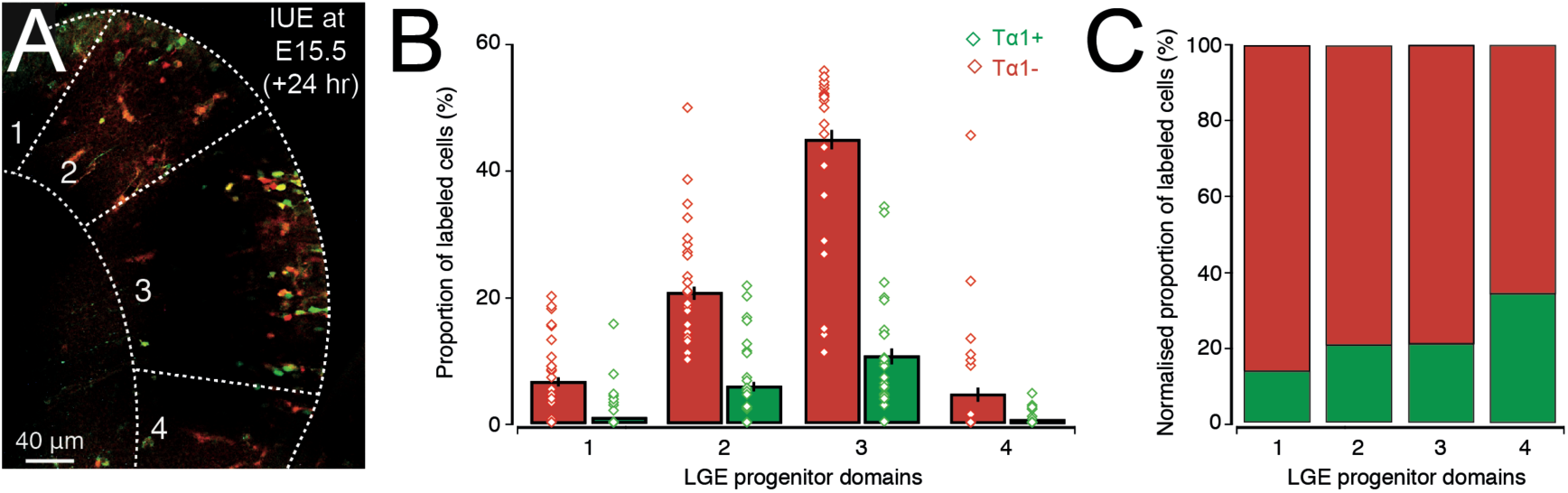
IUE labeled cells can be found in all domains of the LGE. (**A**) Example coronal section of an embryonic mouse brain 24 hours after IUE at E15.5 with Tα1-cre and FLEx plasmids. Note that GFP^+^ and TdTomato^+^ labeled cells can be seen in all progenitor domains of the LGE. (**B**) Average barplot of the proportion of Tα1^+^/GFP^+^ and Tα1^-^/TdTomato^+^ labeled cells in the different progenitor domains of the LGE. Note that our labeling strategy predominantly targets LGE3 (all labeled neurons: LGE1: 7.6%, LGE2: 32.0%, LGE3: 59.0% and LGE4: 1.3%) and that each progenitor domain of the LGE contains both Tα1^+^/GFP^+^ and Tα1^-^ /TdTomato^+^ progenitors. (**C**) The relative proportion of Tα1^+^/GFP^+^ and Tα1^-^ /TdTomato^+^ labeled cells appears relatively constant between the different domains of the LGE.

**Supplementary Figure 2:**
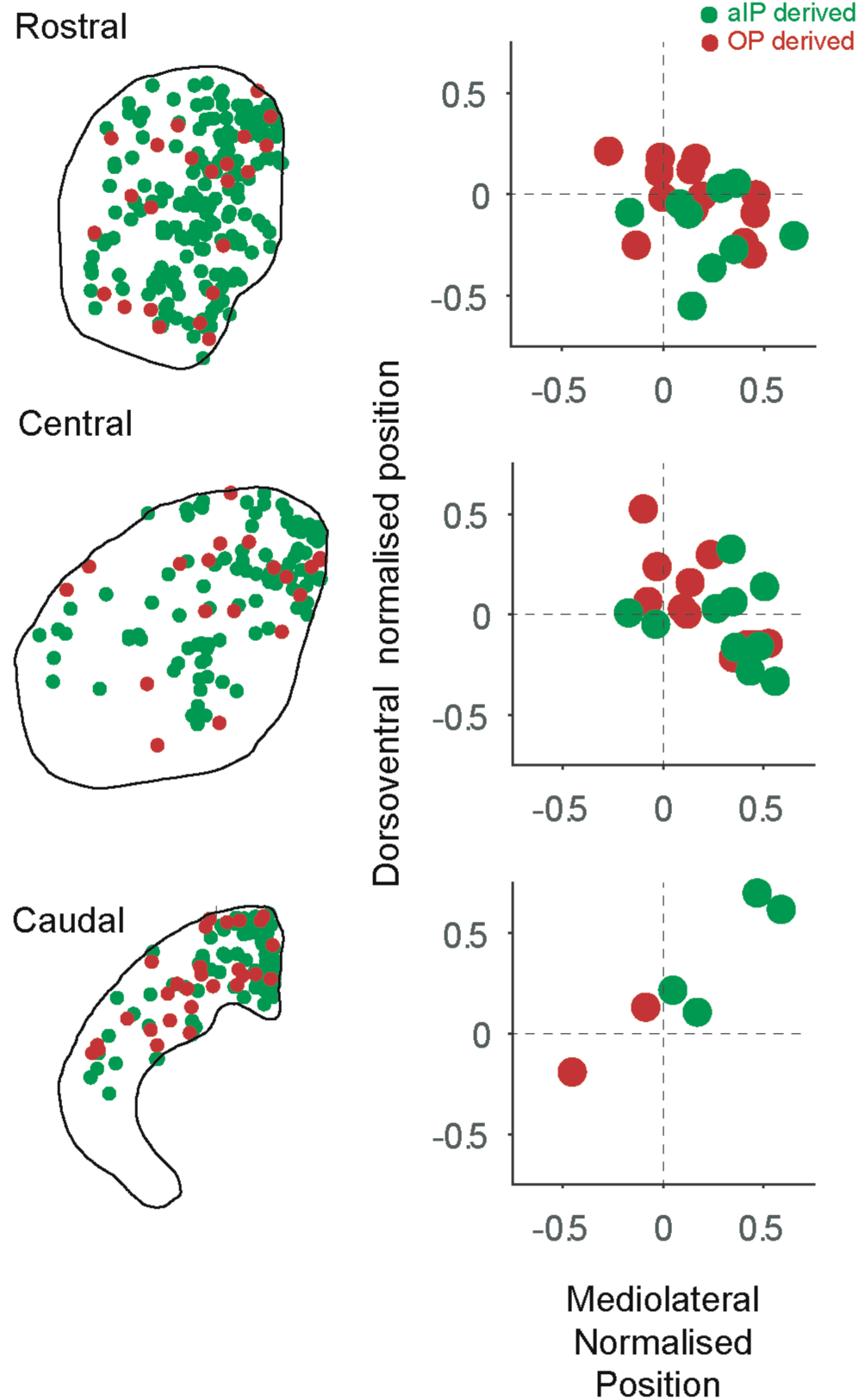
Labeled aIP and OP derived striatal neurons in rostral, central and caudal striatum. Left column, positions of aIP and OP derived neurons in rostral, central and caudal sections of a single animal. Right column, normalised positions of aIP and OP derived neurons in each plane (rostral n = 13, central n = 11, caudal n = 4 mice).

**Supplementary Figure 3:**
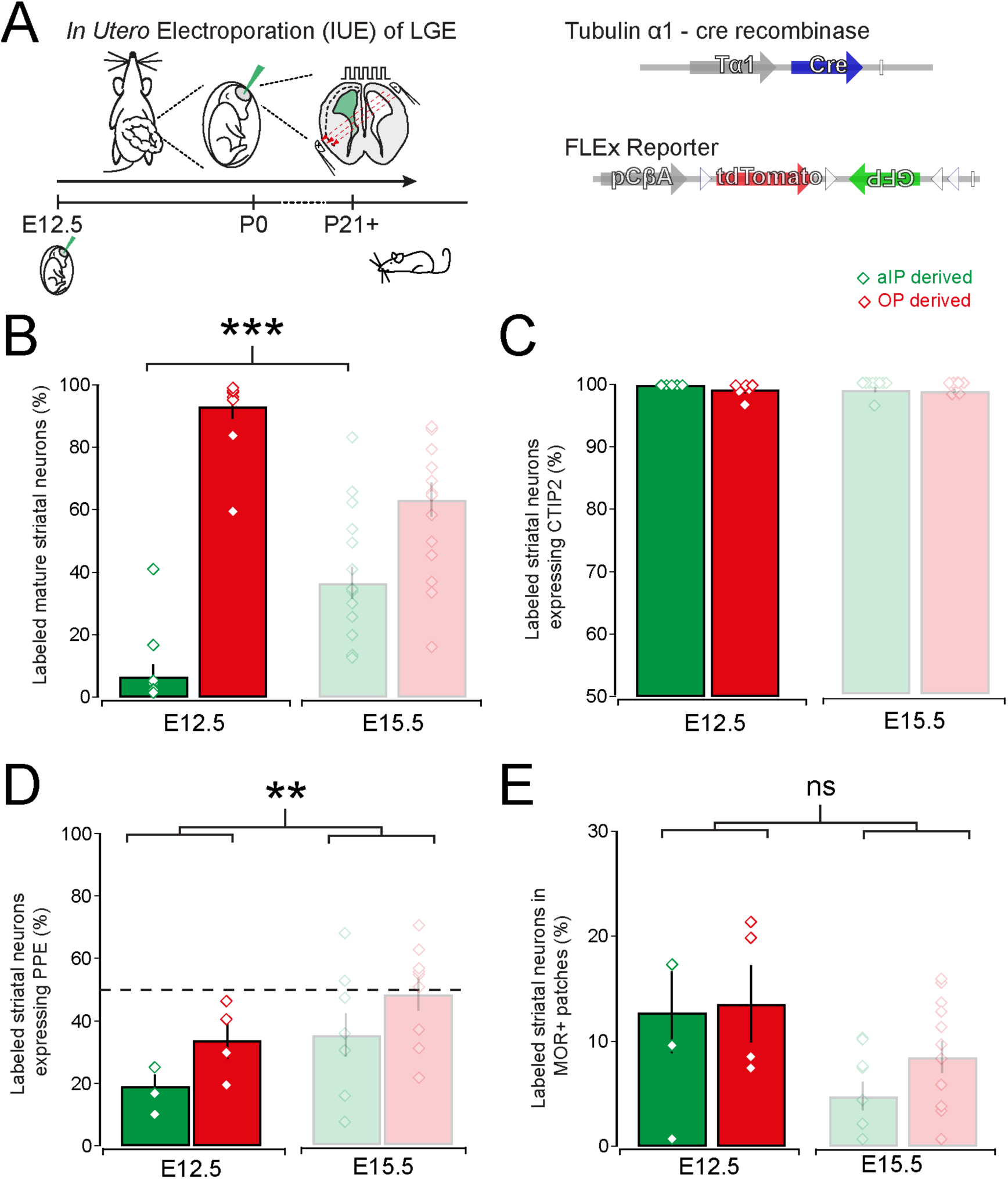
Labeled aIP and OP derived striatal neurons after IUE during early stages of neurogenesis. (**A**) Animals that underwent IUE at E12.5 with Tα1-cre and FLEx reporter plasmids were left to mature untill young adulthood (P21+) (**B**) Significantly more aIP derived neurons are generated using IUE at E15.5 than at E12.5 (E12.5: 6.6 ± 4.0% and E15.5: 36.7 ± 5.5%, Mann-Whitney test, p=0.0002, n = 10 and 16 brains). (**C**) IUE at both E12.5 and E15.5 generate mostly striatal CTIP^+^ neurons. (**D**) There is no significant difference in the production of PPE^+^/D2 SPNs by aIP or OP progenitors at E12.5 or E15.5. However, grouping all labeled neurons together at E12.5 a smaller proportion of PPE^+^/D2 SPNs is labeled at this early stage of neurogenesis (E12.5 vs. E15.5, Mann-Whitney test; p = 0.006; n = 4 and 9 brains). (**E**) There is a trend towards more labeled neurons in the patch compartments after IUE at E12.5 (E12.5: 12.4 ± 2.4% vs. E15.5: 6.9 ± 1.3% in MOR+ patches; Mann-Whitney test, p = 0.170, n = 4 and 12).

**Supplementary Figure 4:**
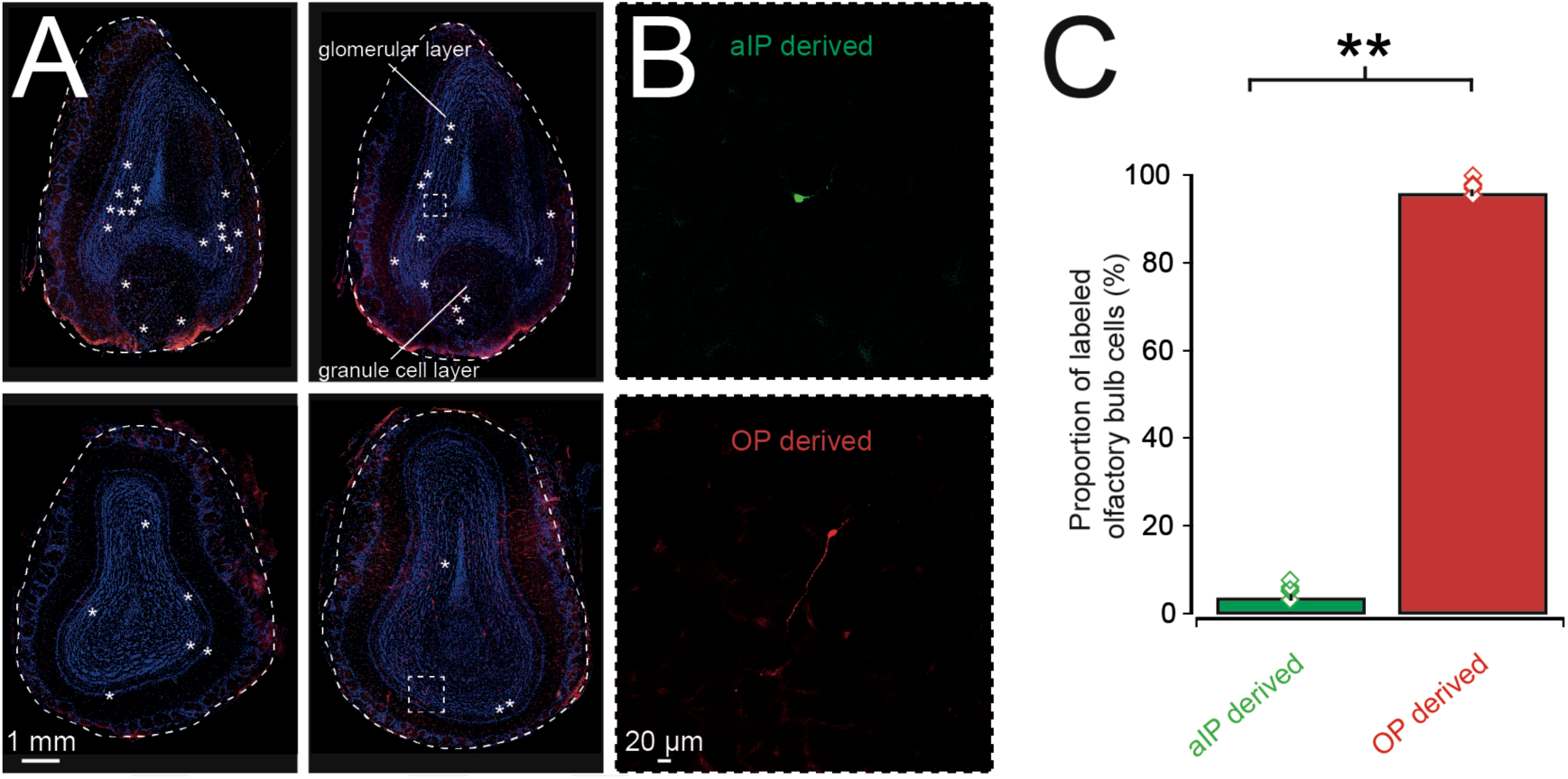
IUE of LGE neural progenitors also labels a small number of olfactory bulb cells. (**A**) Example images of 4 different olfactory bulbs with labeled neurons indicated by asterisks. (**B**) High magnification image of an aIP derived cell (top) and an OP derived cell (bottom) in the olfactory bulb. (**C**) The majority of labeled cells in the olfactory bulb are OP derived (aIP derived: 3.9 ± 1.6% and OP derived: 96.1 ± 1.6%, p=0.0079, Mann-Whitney test, n = 5 brains).

## Notes

**Conflict of interest:** We declare no conflict of interest.

## References

Albin, R.L. (2018) Tourette syndrome: a disorder of the social decision-making network. Brain, 141, 332–347.

Arlotta, P., Molyneaux, B.J., Jabaudon, D., Yoshida, Y. & Macklis, J.D. (2008) Ctip2 controls the differentiation of medium spiny neurons and the establishment of the cellular architecture of the striatum. J Neurosci, 28, 622–632.

Bagetta, V., Picconi, B., Marinucci, S., Sgobio, C., Pendolino, V., Ghiglieri, V., Fusco, F.R., Giampa, C. & Calabresi, P. (2011) Dopamine-dependent long-term depression is expressed in striatal spiny neurons of both direct and indirect pathways: implications for Parkinson’s disease. J Neurosci, 31, 12513–12522.

Baumgart, J. & Baumgart, N. (2016) Cortex-, Hippocampus-, Thalamus-, Hypothalamus-, Lateral Septal Nucleus- and Striatum-specific In Utero Electroporation in the C57BL/6 Mouse. J Vis Exp.

Brand, S. & Rakic, P. (1979) Genesis of the primate neostriatum: [3H]thymidine autoradiographic analysis of the time of neuron origin in the rhesus monkey. Neuroscience, 4, 767–778.

Cadwell, C.R., Scala, F., Fahey, P.G., Kobak, D., Sinz, F.H., Johnsson, P., Li, S., Cotton, R.J., Sandberg, R., Berens, P., Jiang, X. & Tolias, A.S. (2019) Cell type composition and circuit organization of neocortical radial clones. bioRxiv, 526681.

Cepko, C. (2014) Intrinsically different retinal progenitor cells produce specific types of progeny. Nature reviews, 15, 615–627.

Chan, C.S., Peterson, J.D., Gertler, T.S., Glajch, K.E., Quintana, R.E., Cui, Q., Sebel, L.E., Plotkin, J.L., Shen, W., Heiman, M., Heintz, N., Greengard, P. & Surmeier, D.J. (2012) Strain-specific regulation of striatal phenotype in Drd2-eGFP BAC transgenic mice. J Neurosci, 32, 9124–9132.

Chuhma, N., Tanaka, K.F., Hen, R. & Rayport, S. (2011) Functional connectome of the striatal medium spiny neuron. J Neurosci, 31, 1183–1192.

Crittenden, J.R. & Graybiel, A.M. (2011) Basal Ganglia disorders associated with imbalances in the striatal striosome and matrix compartments. Frontiers in neuroanatomy, 5, 59.

Day, M., Wokosin, D., Plotkin, J.L., Tian, X. & Surmeier, D.J. (2008) Differential excitability and modulation of striatal medium spiny neuron dendrites. J Neurosci, 28, 11603–11614.

Del Campo, N., Chamberlain, S.R., Sahakian, B.J. & Robbins, T.W. (2011) The roles of dopamine and noradrenaline in the pathophysiology and treatment of attention-deficit/hyperactivity disorder. Biological psychiatry, 69, e145–157.

Ellender, T.J., Avery, S.V., Mahfooz, K., von Klemperer, A., Nixon, S.L., Buchan, M.J., van Rheede, J.J., Gatti, A., Waites, C., Newey, S.E. & Akerman, C.J. (2018) Fine-scale excitatory cortical circuits reflect embryonic progenitor pools. bioRxiv.

Fishell, G., Rossant, J. & van der Kooy, D. (1990) Neuronal lineages in chimeric mouse forebrain are segregated between compartments and in the rostrocaudal and radial planes. Dev Biol, 141, 70–83.

Flames, N., Pla, R., Gelman, D.M., Rubenstein, J.L., Puelles, L. & Marin, O. (2007) Delineation of multiple subpallial progenitor domains by the combinatorial expression of transcriptional codes. J Neurosci, 27, 9682–9695.

Franco, S.J., Gil-Sanz, C., Martinez-Garay, I., Espinosa, A., Harkins-Perry, S.R., Ramos, C. & Muller, U. (2012) Fate-Restricted Neural Progenitors in the Mammalian Cerebral Cortex. Science, 337, 746–749.

Franco, S.J. & Muller, U. (2013) Shaping our minds: stem and progenitor cell diversity in the mammalian neocortex. Neuron, 77, 19–34.

Franklin, K.B.J. & Paxinos, G. (2008) The mouse brain in stereotaxic coordinates. Elsevier Academic Press, Amsterdam; London.

Gal, J.S., Morozov, Y.M., Ayoub, A.E., Chatterjee, M., Rakic, P. & Haydar, T.F. (2006) Molecular and morphological heterogeneity of neural precursors in the mouse neocortical proliferative zones. J Neurosci, 26, 1045–1056.

Garas, F.N., Kormann, E., Shah, R.S., Vinciati, F., Smith, Y., Magill, P.J. & Sharott, A. (2018) Structural and molecular heterogeneity of calretinin-expressing interneurons in the rodent and primate striatum. The Journal of comparative neurology, 526, 877–898.

Garas, F.N., Shah, R.S., Kormann, E., Doig, N.M., Vinciati, F., Nakamura, K.C., Dorst, M.C., Smith, Y., Magill, P.J. & Sharott, A. (2016) Secretagogin expression delineates functionally-specialized populations of striatal parvalbumin-containing interneurons. Elife, 5.

Gerfen, C.R. (1984) The neostriatal mosaic: compartmentalization of corticostriatal input and striatonigral output systems. Nature, 311, 461–464.

Gerfen, C.R. (1992) The neostriatal mosaic: multiple levels of compartmental organization. Trends in neurosciences, 15, 133–139.

Gerfen, C.R., Baimbridge, K.G. & Miller, J.J. (1985) The neostriatal mosaic: compartmental distribution of calcium-binding protein and parvalbumin in the basal ganglia of the rat and monkey. Proceedings of the National Academy of Sciences of the United States of America, 82, 8780–8784.

Gerfen, C.R., Engber, T.M., Mahan, L.C., Susel, Z., Chase, T.N., Monsma, F.J., Jr. & Sibley, D.R. (1990) D1 and D2 dopamine receptor-regulated gene expression of striatonigral and striatopallidal neurons. Science, 250, 1429–1432.

Gertler, T.S., Chan, C.S. & Surmeier, D.J. (2008) Dichotomous anatomical properties of adult striatal medium spiny neurons. J Neurosci, 28, 10814–10824.

Gong, S., Zheng, C., Doughty, M.L., Losos, K., Didkovsky, N., Schambra, U.B., Nowak, N.J., Joyner, A., Leblanc, G., Hatten, M.E. & Heintz, N. (2003) A gene expression atlas of the central nervous system based on bacterial artificial chromosomes. Nature, 425, 917–925.

Graybiel, A.M. & Hickey, T.L. (1982) Chemospecificity of ontogenetic units in the striatum: demonstration by combining [3H]thymidine neuronography and histochemical staining. Proceedings of the National Academy of Sciences of the United States of America, 79, 198–202.

Graybiel, A.M. & Ragsdale, C.W., Jr. (1978) Histochemically distinct compartments in the striatum of human, monkeys, and cat demonstrated by acetylthiocholinesterase staining. Proceedings of the National Academy of Sciences of the United States of America, 75, 5723–5726.

Graybiel, A.M. & Rauch, S.L. (2000) Toward a neurobiology of obsessive-compulsive disorder. Neuron, 28, 343–347.

Grillner, S. & Robertson, B. (2016) The Basal Ganglia Over 500 Million Years. Current biology : CB, 26, R1088–R1100.

Guillamon-Vivancos, T., Tyler, W.A., Medalla, M., Chang, W.W., Okamoto, M., Haydar, T.F. & Luebke, J.I. (2019) Distinct Neocortical Progenitor Lineages Fine-tune Neuronal Diversity in a Layer-specific Manner. Cereb Cortex, 29, 1121–1138.

Guo, Q., Wang, D., He, X., Feng, Q., Lin, R., Xu, F., Fu, L. & Luo, M. (2015) Whole-brain mapping of inputs to projection neurons and cholinergic interneurons in the dorsal striatum. PLoS ONE, 10, e0123381.

Halliday, A.L. & Cepko, C.L. (1992) Generation and migration of cells in the developing striatum. Neuron, 9, 15–26.

Herkenham, M., Edley, S.M. & Stuart, J. (1984) Cell clusters in the nucleus accumbens of the rat, and the mosaic relationship of opiate receptors, acetylcholinesterase and subcortical afferent terminations. Neuroscience, 11, 561–593.

Hunnicutt, B.J., Jongbloets, B.C., Birdsong, W.T., Gertz, K.J., Zhong, H. & Mao, T. (2016) A comprehensive excitatory input map of the striatum reveals novel functional organization. Elife, 5.

Jimenez-Castellanos, J. & Graybiel, A.M. (1989) Compartmental origins of striatal efferent projections in the cat. Neuroscience, 32, 297–321.

Kelly, S.M., Raudales, R., He, M., Lee, J.H., Kim, Y., Gibb, L.G., Wu, P., Matho, K., Osten, P., Graybiel, A.M. & Huang, Z.J. (2018) Radial Glial Lineage Progression and Differential Intermediate Progenitor Amplification Underlie Striatal Compartments and Circuit Organization. Neuron, 99, 345–361 e344.

Kowalczyk, T., Pontious, A., Englund, C., Daza, R.A., Bedogni, F., Hodge, R., Attardo, A., Bell, C., Huttner, W.B. & Hevner, R.F. (2009) Intermediate neuronal progenitors (basal progenitors) produce pyramidal-projection neurons for all layers of cerebral cortex. Cereb Cortex, 19, 2439–2450.

Krajeski, R.N., Macey-Dare, A., van Heusden, F., Ebrahimjee, F. & Ellender, T.J. (2018) Early postnatal development of the cellular and circuit properties of striatal D1 and D2 spiny projection neurons. bioRxiv.

Kramer, P.F., Christensen, C.H., Hazelwood, L.A., Dobi, A., Bock, R., Sibley, D.R., Mateo, Y. & Alvarez, V.A. (2011) Dopamine D2 receptor overexpression alters behavior and physiology in Drd2-EGFP mice. J Neurosci, 31, 126–132.

Kriegstein, A. & Alvarez-Buylla, A. (2009) The glial nature of embryonic and adult neural stem cells. Annual review of neuroscience, 32, 149–184.

Langen, M., Kas, M.J., Staal, W.G., van Engeland, H. & Durston, S. (2011) The neurobiology of repetitive behavior: of mice. Neurosci Biobehav Rev, 35, 345–355.

Lee, T., Kaneko, T., Taki, K. & Mizuno, N. (1997) Preprodynorphin-, preproenkephalin-, and preprotachykinin-expressing neurons in the rat neostriatum: an analysis by immunocytochemistry and retrograde tracing. The Journal of comparative neurology, 386, 229–244.

Lobo, M.K., Karsten, S.L., Gray, M., Geschwind, D.H. & Yang, X.W. (2006) FACS-array profiling of striatal projection neuron subtypes in juvenile and adult mouse brains. Nature neuroscience, 9, 443–452.

Marchand, R. & Lajoie, L. (1986) Histogenesis of the striopallidal system in the rat. Neurogenesis of its neurons. Neuroscience, 17, 573–590.

Mason, H.A., Rakowiecki, S.M., Raftopoulou, M., Nery, S., Huang, Y., Gridley, T. & Fishell, G. (2005) Notch signaling coordinates the patterning of striatal compartments. Development, 132, 4247–4258.

McNaught, K.S. & Mink, J.W. (2011) Advances in understanding and treatment of Tourette syndrome. Nat Rev Neurol, 7, 667–676.

Nelson, A.B., Hang, G.B., Grueter, B.A., Pascoli, V., Luscher, C., Malenka, R.C. & Kreitzer, A.C. (2012) A comparison of striatal-dependent behaviors in wild-type and hemizygous Drd1a and Drd2 BAC transgenic mice. J Neurosci, 32, 9119–9123.

Newman, H., Liu, F.C. & Graybiel, A.M. (2015) Dynamic ordering of early generated striatal cells destined to form the striosomal compartment of the striatum. The Journal of comparative neurology, 523, 943–962.

Noctor, S.C., Flint, A.C., Weissman, T.A., Dammerman, R.S. & Kriegstein, A.R. (2001) Neurons derived from radial glial cells establish radial units in neocortex. Nature, 409, 714–720.

Noctor, S.C., Martinez-Cerdeno, V., Ivic, L. & Kriegstein, A.R. (2004) Cortical neurons arise in symmetric and asymmetric division zones and migrate through specific phases. Nature neuroscience, 7, 136–144.

Olsson, M., Bjorklund, A. & Campbell, K. (1998) Early specification of striatal projection neurons and interneuronal subtypes in the lateral and medial ganglionic eminence. Neuroscience, 84, 867–876.

Pan, W.X., Mao, T. & Dudman, J.T. (2010) Inputs to the dorsal striatum of the mouse reflect the parallel circuit architecture of the forebrain. Frontiers in neuroanatomy, 4, 147.

Pert, C.B., Kuhar, M.J. & Snyder, S.H. (1976) Opiate receptor: autoradiographic localization in rat brain. Proceedings of the National Academy of Sciences of the United States of America, 73, 3729–3733.

Pilz, G.A., Shitamukai, A., Reillo, I., Pacary, E., Schwausch, J., Stahl, R., Ninkovic, J., Snippert, H.J., Clevers, H., Godinho, L., Guillemot, F., Borrell, V., Matsuzaki, F. & Gotz, M. (2013) Amplification of progenitors in the mammalian telencephalon includes a new radial glial cell type. Nature communications, 4, 2125.

Planert, H., Szydlowski, S.N., Hjorth, J.J., Grillner, S. & Silberberg, G. (2010) Dynamics of synaptic transmission between fast-spiking interneurons and striatal projection neurons of the direct and indirect pathways. J Neurosci, 30, 3499–3507.

Reid, C.B. & Walsh, C.A. (2002) Evidence of common progenitors and patterns of dispersion in rat striatum and cerebral cortex. J Neurosci, 22, 4002–4014.

Saunders, A., Johnson, C.A. & Sabatini, B.L. (2012) Novel recombinant adeno-associated viruses for Cre activated and inactivated transgene expression in neurons. Frontiers in neural circuits, 6, 47.

Schambra, U.B. & Schambra, U.B.A.o.p.m.b. (2008) Prenatal mouse brain atlas. Springer, New York; London.

Sharott, A., Vinciati, F., Nakamura, K.C. & Magill, P.J. (2017) A Population of Indirect Pathway Striatal Projection Neurons Is Selectively Entrained to Parkinsonian Beta Oscillations. J Neurosci, 37, 9977–9998.

Shepherd, G.M. (2013) Corticostriatal connectivity and its role in disease. Nature reviews, 14, 278–291.

Shitamukai, A., Konno, D. & Matsuzaki, F. (2011) Oblique radial glial divisions in the developing mouse neocortex induce self-renewing progenitors outside the germinal zone that resemble primate outer subventricular zone progenitors. J Neurosci, 31, 3683–3695.

Smart, I.H. (1976) A pilot study of cell production by the ganglionic eminences of the developing mouse brain. J Anat, 121, 71–84.

Stancik, E.K., Navarro-Quiroga, I., Sellke, R. & Haydar, T.F. (2010) Heterogeneity in ventricular zone neural precursors contributes to neuronal fate diversity in the postnatal neocortex. J Neurosci, 30, 7028–7036.

Stenman, J., Toresson, H. & Campbell, K. (2003) Identification of two distinct progenitor populations in the lateral ganglionic eminence: implications for striatal and olfactory bulb neurogenesis. J Neurosci, 23, 167–174.

Stopczynski, R.E., Poloskey, S.L. & Haber, S.N. (2008) Cell proliferation in the striatum during postnatal development: preferential distribution in subregions of the ventral striatum. Brain structure & function, 213, 119–127.

Tan, S.S. & Breen, S. (1993) Radial mosaicism and tangential cell dispersion both contribute to mouse neocortical development. Nature, 362, 638–640.

Taverna, E., Gotz, M. & Huttner, W.B. (2014) The cell biology of neurogenesis: toward an understanding of the development and evolution of the neocortex. Annu Rev Cell Dev Biol, 30, 465–502.

Taverna, S., Ilijic, E. & Surmeier, D.J. (2008) Recurrent collateral connections of striatal medium spiny neurons are disrupted in models of Parkinson’s disease. J Neurosci, 28, 5504–5512.

Tinterri, A., Menardy, F., Diana, M.A., Lokmane, L., Keita, M., Coulpier, F., Lemoine, S., Mailhes, C., Mathieu, B., Merchan-Sala, P., Campbell, K., Gyory, I., Grosschedl, R., Popa, D. & Garel, S. (2018) Active intermixing of indirect and direct neurons builds the striatal mosaic. Nature communications, 9, 4725.

Tyler, W.A., Medalla, M., Guillamon-Vivancos, T., Luebke, J.I. & Haydar, T.F. (2015) Neural precursor lineages specify distinct neocortical pyramidal neuron types. J Neurosci, 35, 6142–6152.

van der Kooy, D. & Fishell, G. (1987) Neuronal birthdate underlies the development of striatal compartments. Brain research, 401, 155–161.

Wamsley, B. & Fishell, G. (2017) Genetic and activity-dependent mechanisms underlying interneuron diversity. Nature reviews, 18, 299–309.

Wang, X., Tsai, J.W., LaMonica, B. & Kriegstein, A.R. (2011) A new subtype of progenitor cell in the mouse embryonic neocortex. Nature neuroscience, 14, 555–561.

Wonders, C.P. & Anderson, S.A. (2006) The origin and specification of cortical interneurons. Nature reviews, 7, 687–696.

Xu, Z., Liang, Q., Song, X., Zhang, Z., Lindtner, S., Li, Z., Wen, Y., Liu, G., Guo, T., Qi, D., Wang, M., Wang, C., Li, H., You, Y., Wang, X., Chen, B., Feng, H., Rubenstein, J.L. & Yang, Z. (2018) SP8 and SP9 coordinately promote D2-type medium spiny neuron production by activating Six3 expression. Development, 145.

Yu, Y.C., Bultje, R.S., Wang, X. & Shi, S.H. (2009) Specific synapses develop preferentially among sister excitatory neurons in the neocortex. Nature, 458, 501–504.

Yu, Y.C., He, S., Chen, S., Fu, Y., Brown, K.N., Yao, X.H., Ma, J., Gao, K.P., Sosinsky, G.E., Huang, K. & Shi, S.H. (2012) Preferential electrical coupling regulates neocortical lineage-dependent microcircuit assembly. Nature, 486, 113–117.

